# Morphine Re-arranges Chromatin Spatial Architecture of Primate Cortical Neurons

**DOI:** 10.1101/2023.03.06.531278

**Authors:** Liang Wang, Xiaojie Wang, Chunqi Liu, Wei Xu, Weihong Kuang, Qian Bu, Hongchun Li, Ying Zhao, Linhong Jiang, Yaxing Chen, Feng Qin, Shu Li, Qinfan Wei, Xiaocong Liu, Bin Liu, Yuanyuan Chen, Yanping Dai, Hongbo Wang, Jingwei Tian, Gang Cao, Yinglan Zhao, Xiaobo Cen

**Author notes:** Corresponding author. E-mail address (Cen X). Equal contribution (Liang Wang, Xiaojie Wang, Chunqi Liu).

## Abstract

The expression of linear DNA sequences is precisely regulated by the three–dimensional (3D) architecture of chromatin. Morphine-induced aberrant gene networks of neurons have been extensively investigated; however, how morphine impacts the 3D genomic architecture of neuorns is still unknown. Here, we applied digestion-ligation-only high-throughput chromosome conformation capture (DLO Hi-C) technology to investigate the affection of morphine on 3D chromatin architecture of primate cortical neurons. After receiving continuous morphine administration for 90 days on rhesus monkeys, we discovered that morphine re-arranged chromosome territories, with a total of 391 segmented compartments being switched. Morphine altered over half of the detected topologically associated domains (TADs), most of which exhibited a variety of shifts, followed by separating and fusing types. Analysis of the looping events at kilobase-scale resolution revealed that morphine increased not only the number but also the length of differential loops. Moreover, all identified differentially expressed genes (DEGs) from the RNA sequencing (RNA-seq) were mapped to the specific TAD boundaries or differential loops, and were further validated to be significantly changed. Collectively, an altered 3D genomic architecture of cortical neurons may regulate the gene networks associated-morphine effects. Our finding provides critical hubs connecting chromosome spatial organization and gene networks associated with the morphine effects in humans.

## Introduction

Morphine, the most effective opioid analgesic, is widely used in clinics for the management of chronic and severe acute pain. However, long-term morphine administration induces tolerance and concomitant hyperalgesia, which severely limits its efficacy and application in clinics. Current knowledge on morphine-induced psychiatric behaviors and alteration of gene expression profile in the brain mostly emphasizes the neuroadaptive changes in neural plasticity and circuit, synaptic receptor desensitization, and neurotransmitter release [1,2]. An important mechanism underlying this complex neuronal malfunction is gene misexpression in the central nervous system (CNS), which is orchestrated by a network of transcription factors and chromatin–remodeling enzymes [3,4]. Long-term or even a single morphine injection remarkably dysregulates the expressions of a cluster of critical neuronal genes, such as *c-fos* [4], *Oprm1* [5], *Arc*, *BDNF*, and *NGF* [6]. Despite these advances, the basic folding principles of the sophisticated effects of morphine-modified epigenome have not been revealed, and the role of chromatin structural changes in the pharmacological and toxicological effects of morphine remains unknown.

Gene expression is precisely controlled by proper folding of chromatin structure, a representative functional unit of the genome, enabling distal regulatory elements to regulate the expression of target genes even megabase (Mb) away at the linear genome maps [7]. Perturbation of three–dimensional (3D) chromatin architecture is a cause of gene misexpression, which contributes to a variety of human illnesses and developmental disorders [8–11]. For instance, an architecture variation at *Sox9* loci causes the incorporation of a neighboring *Kcnj2* gene in another neo-topologically associated domain (TAD), which subsequently induces ectopic contacts of *Kcnj2* with the regulatory elements, eventually leading to a limb malformation [11]. In 3D chromatin architecture, chromatin DNA together with structural proteins is hierarchically packaged into a multi-layered spatial structure, from loops to TADs, compartmentalized structures, and chromosome territories [12].

Previous studies have shown that epigenetic changes, such as increased histone acetylation and DNA methylation, play a critical role in mediating morphine effects [13,14]. However, it is still unknown how the 3D configuration of the genome is linked to morphine effects. The importance of accurate detection and interpretation of large-scale genomic rearrangements is highlighted by the fact that chromatin conformation frequently adjusts its high-order structure to accommodate the different biological processes [15]. Recent advances in chromosome conformation capture technology have greatly broadened an insight into 3D chromatin spatial architecture. Compared to 3C technology, high-throughput chromosome conformation capture (Hi-C) technology allows for simultaneous interrogation of all contact loci, resulting in a comprehensive visualization of all–to–all genome–wide interactions with unprecedented resolution in combination with high-throughput sequencing [2,10,16,17].

3D chromatin architecture is not particularly conserved with less than 30% TAD sharing across species [18], and the significance of chromatin organization is highlighted in the evolution of gene regulation across different lineages. In the present study, through a combination of digestion–ligation–only (DLO) Hi-C technology [19] and genome–wide RNA sequencing (RNA-seq), we investigated the impact of morphine on the chromatin architecture and transcriptional profile of genes of cortical neurons in the rhesus monkey (*Macaca mulatta*), a non–human primate with high genetic similarities to the human genome. Our finding revealed a specific chromatin spatial organization and alteration at different hierarchical levels of cortical neurons after long-term morphine exposure, which bridges the gap between genomic architecture and morphine-modified gene networks.

## Results

### Long-term morphine administration modulates chromatin spatial organization of cortical neurons in the non–human primates

Transcriptional responses of neurons in response to morphine have been extensively investigated; however, the alteration of genome 3D architecture is still unknown. We applied DLO Hi-C technology to investigate the alteration of DNA spatial structure of cortical neurons in rhesus monkeys treated with morphine for 3 months continuously [19]. To reduce the genetic differences, four male rhesus monkeys born from two fathers were selected for this study. Each pair of male monkeys from the same father was divided into saline and morphine groups (n = 2/group). Two monkeys in the morphine group were injected subcutaneously with morphine three times a day for 90 days continuously, with a cumulative dose regimen of 3, 6, 9, and 12 mg/kg for the first four weeks, respectively, and a constant dosage of 15 mg/kg for the remaining days (**Figure 1A**) [2]. As a mock control, the other two rhesus monkeys in the saline group were subcutaneously injected three times a day with the same volume of 0.9% saline for 90 days continuously. The bodyweight of all monkeys was scaled weekly, and the results showed that morphine-treated monkeys exhibited weak retardation in weight gain, whereas saline-treated monkeys grew normally (Figure S1A). Fourteen hours after the last injection, the spontaneous withdrawal signs were individually monitored and scored at five consecutive periods: 0.5, 1, 1.5, 2, and 2.5 h, respectively. We found that the cumulative number of withdrawal signs was significantly higher in the morphine group than in the saline group (Figure 1B and Table S1). Within an observation period of 2.5 h, in contrast to 6.5 of total withdrawal signs in the saline group, the withdrawal scores accumulated to 37.5 in the morphine group, indicating an obvious tolerance and physical dependence after long-term morphine administration. In detail, the increased withdrawal scores were attributed to diverse withdrawal signs, including vocalizing, tremors, restlessness, lying on the side or abdomen, and fighting, with a total score of 7.5, 7, 4.5, 3.5, and 3.5, respectively (Table S1). These withdrawal signs were similar to those characteristic symptoms of opioid withdrawal in humans [16,20].

**Figure 1.**
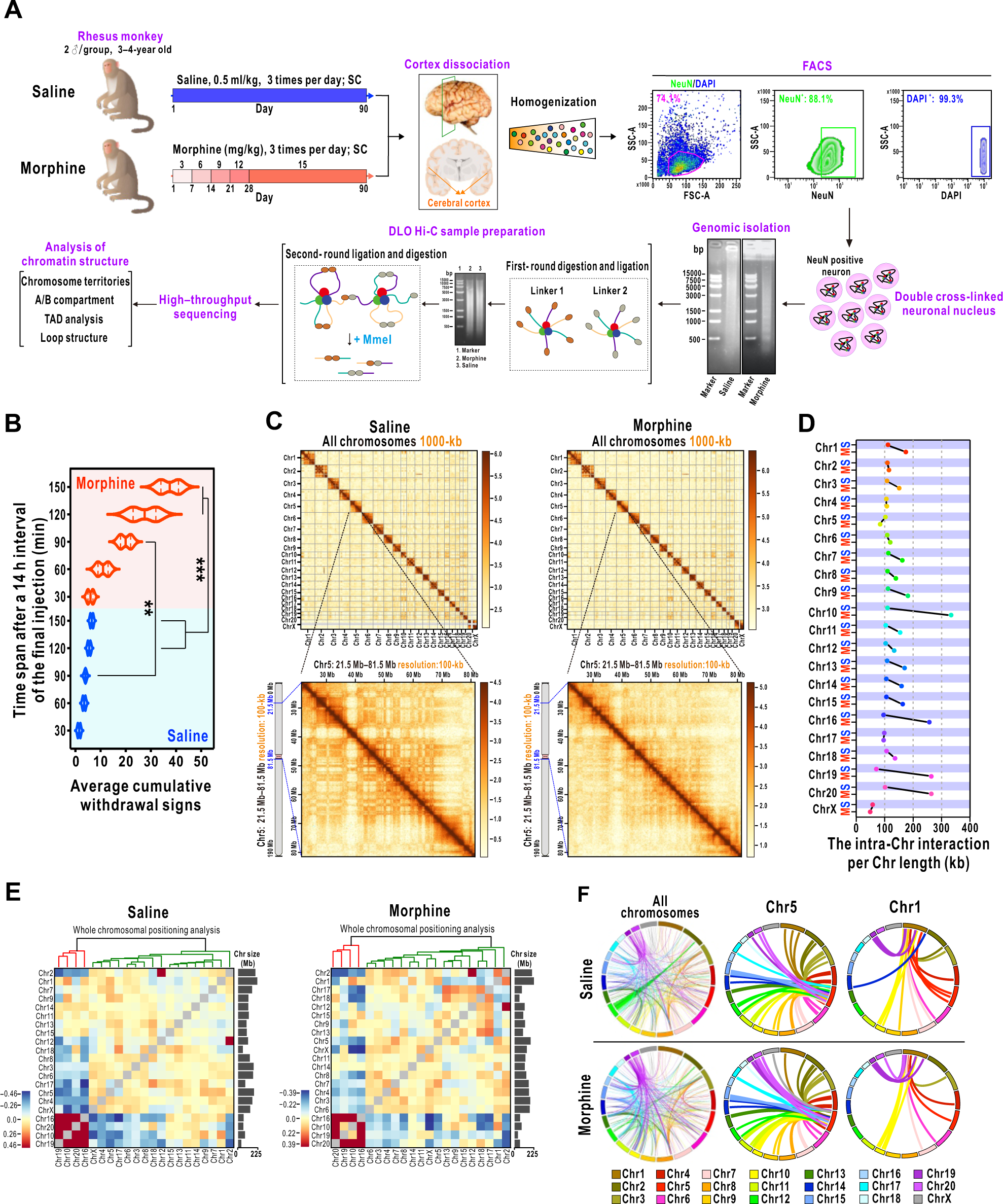
Morphine re-arranges the chromatin spatial architecture of non-human primate cortical neurons. **A.** Schematic workflow of experiment procedure. **B.** Violin plot shows cumulative scores of spontaneous withdrawal signs observed in monkeys during a period of 30, 60, 90, 120, and 150 min, respectively, 14 h after the final injection. Two-way ANOVA, \*\**P* < 0.01, \*\*\**P* < 0.001. **C.** Heatmaps show normalized DLO Hi-C interaction frequencies at different resolutions. 1000 kb for the entire genome and 100 kb for the selected area of Chr5 (21.5 Mb–81.5 Mb). **D.** The intra-Chr interaction frequencies per Chr length (kb). The x-axis represents the valid paired reads of intra-chromosomal interaction by dividing the full length of each chromosome (kb). S: saline, M: morphine. **E.** Heatmaps of chromosome positioning sorting by hierarchically clustered across the entire genome. **F.** Circos plots of the first 1000 inter-chromosomal interactions across the entire genome, and the trans-interaction of Chr5 and Chr1 with the other chromosomes. DLO Hi-C, digestion–ligation–only high–throughput chromosome conformation capture. Intra-Chr, intra-chromosomal, Chr, chromosome.

Next, the cerebral cortex was dissected for exploring the 3D chromatin organization as well as differences in genome-wide transcriptional between the two groups. The fresh cortex was enzymatically dissociated, and subsequently double-labeled with nuclear marker 4′,6-diamidino-2-phenylindole (DAPI) and the mature neuronal marker neuronal nuclei (NeuN). The NeuN^+^ and DAPI^+^ neurons were sorted through a fluorescence–activated–cell–sorting (FACS)-based isolation technique (Figure 1A and Figure S1B). The cross-linked chromatin was extracted from isolated cortical neurons and subjected to DLO Hi-C analysis as previously reported [19] (Figure 1A). The high-throughput sequencing yielded more than 850 million raw reads for each condition. The mappability of reads was 60.91% and 51.61%, and the overall inter- and intra-chromosomal interaction ratios were 6.62% and 21.58% in the saline group, and 5.95% and 17.72% in the morphine group, respectively (Figure S2A). We surveyed 3D genome organization and analyzed features across several scales, including chromatin territories, compartments, TADs, and loops. The interaction matrices of the whole genome presented similar interaction patterns between the two groups at the entire genome level (Figure 1C and Figure S2B). Consistently, when interaction frequencies were plotted as a function of the genomic distance between loci, the contact frequency of whole or individual chromosomes was identical in both groups, with the exception of a modest increase around 100 Mb distance in the morphine group (Figures S2C and S2D).

Even though the contact maps of the whole genome were only slightly affected, morphine treatment resulted in evident modifications in genome structure at different chromosomal scales. For example, the representative cis-interaction matrices of Chr5 observed at different resolutions (1000-kb, 100-kb, and 10-kb resolution, respectively) exhibited a reduced intra-chromosomal contact frequency in the morphine group (Figure 1C and Figure S2B). The intra-chromosomal interaction frequency of all chromosomes was further quantified and compared. Except for Chr5, 17, and X, the intra-chromosomal interaction of most chromosomes was clearly increased by morphine after normalization to the respective chromosome length (kb) (Figure 1D and Table S2). Collectively, these data indicated that morphine may alter the chromatin 3D spatial architecture of cortical neurons.

### Morphine re-arranges the chromosome territories of cortical neurons

The frequency of genomic contact, including both inter- and intra-chromosomal interaction, is altered with the re-arranged chromosome territories [21,22]. We asked whether the varied contact frequencies of each chromosome were attributed to the changed chromosome territories. To this end, the interaction map of all chromosomes was clustered based on the interaction frequencies of each chromosome with the other chromosomes (Figure 1E). After quantifying both the inter- and intra-chromosomal interaction of two conditions, we observed a reduced inter-chromosomal interaction ratio in six chromosomes and an increased interaction ratio in the rest of the chromosomes after morphine administration (Figure S2E). In contrast to the other chromosomes, the inter-chromosomal interaction ratio of the two longest chromosomes, Chr1 and Chr2, was barely affected, indicating a relatively steady state of chromosome territory in response to morphine. Different from Chr1 and Chr2, the inter-chromosomal interaction of Chr5 and Chr19 was significantly altered by morphine, indicating that the territory of these two chromosomes was evidently altered by morphine (Figure S2E and Table S3). Overall, these alterations resulted in a higher inter-chromosomal interaction ratio across the entire genome in the morphine group (Figure S2F).

We further visualized the first 1000 inter-chromosomal interactions of all chromosomes as well as the individual chromosome. By contrast, the overall inter-chromosomal interaction profile was clearly altered by morphine (Figure 1F, left panel). Among all chromosomes, morphine increased the frequency of Chr5 interacting with the other chromosomes, while it lowered the frequency of Chr1 and Chr9 interacting with the other chromosomes (Figure 1F and Figure S3). These findings suggested that morphine can re-arrange chromosome territories of cortical neurons.

### Morphine attenuates genome compartmentalization of cortical neurons

Given the changes in chromosome territories, we next asked whether the compartment status of 3D organization was altered by morphine. Firstly, the eigenvectors retrieved the top three principal components (PC) (PC1, PC2, and PC3) of each chromosome were calculated, and the absolute value was shown in **Figure 2A**. The eigenvector aligning with the largest absolute value, most GC content or gene density was applied to define the A/B compartment profile, with regions of the positive and negative value corresponding, respectively, to A-type (red) and B-type (blue) compartments. The active A and inactive B compartment patterns at the chromosomal scale were individually examined (Figure 2B, Tables S4, and S5). The representative compartment pattern as exemplified by the changes was visualized on Chr5 (Figure 2C). We found that a plaid pattern involving long-range *cis* contacts in Hi-C contact matrices of the saline group was weakened by morphine, suggesting high variability in long-range genomic interactions.

**Figure 2.**
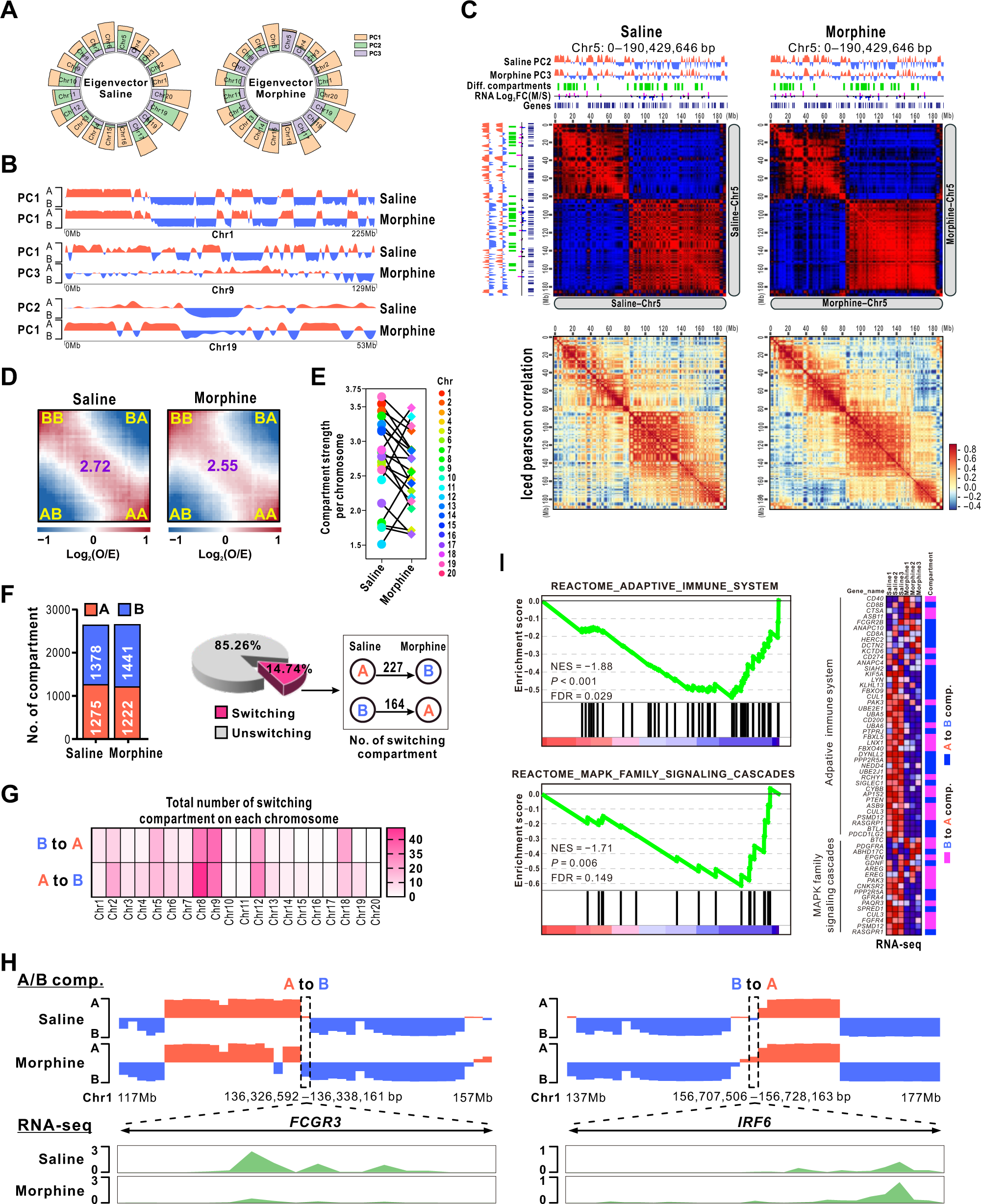
Morphine attenuates spatial compartmentalization of the genome. **A.** Radial stacked plots present the absolute value of the top three eigenvectors of individual chromosomes. PC1, light orange; PC2, light green; and PC3, light purple. **B.** The A-type (red) and B-type (blue) compartment status of Chr1, 9, and 19 at the chromosomal scale using the largest absolute value of the eigenvector. **C.** Compartment weakening for Chr5 is shown in morphine versus saline (upper lane). The iced Pearson correlation of interaction at Chr5 is presented at the bottom. Top and left: the compartment status of saline and morphine are visualized through the PC2 and PC3, respectively. Compartments A and B are separately presented in red and blue color, and the switching regions are the differential (diff.) compartments that are shown in the green line. The significantly up- and down-regulated genes are shown in magenta and blue. The distribution of exons is shown in the dark blue line. **D.** Compartmentalization saddle plots of intra-chromosomal interaction frequencies binned at 1000 kb resolution. The number in the middle represents the overall strength of compartmentalization. **E.** The line graph shows the compartmentalization strength of each chromosome. **F.** Bar graph presents the number of compartments A (red) and B (blue) in two conditions (left panel). The proportion of switching and unswitching compartments between the two groups is displayed in the pie chart (middle panel). The total number of switching compartments is presented in the right panel. **G.** Heatmap displays the number of switching compartments on each chromosome. **H.** Examples of regions in chr1 that switch from A to B (left panel) and B to A (right panel). RNA-seq peaks mapping to significantly changed genes within the switching region are shown in the lower panel. **I.** Enrichment plots enriched in the GSEA Reactome subset. The profiles present the running enrichment score and positions of gene set members on the rank-ordered list (left panel). A Heatmap of the total core genes is shown in the right panel. The expression value of each gene was represented as colors ranging from red (high expression), pink (moderate), light blue (low) to dark blue (lowest expression). The compartment status of each gene following morphine treatment was shown in magenta (compartment B to A) and blue (compartment A to B), respectively. PC, principal component; GSEA, gene set enrichment analysis, NES, normalized enrichment score; FDR, false discovery rate.

We next examined compartment segregation by quantifying compartment strength and found that morphine caused weakened compartmentalization (Figure 2D). The genome-wide compartment strength was attenuated from 2.72 in the saline group to 2.55 in the morphine group. The attenuated compartment strength was attributed to the reduction of all chromosomes with exception of four chromosomes (Chr 7, 11, 12, and 19) (Figure 2E). This data suggested that morphine treatment might diminish the segregation of A and B compartments across the entire genome. Moreover, the number of A/B compartments was also affected by morphine. In the saline and morphine groups, 1275 and 1222 of A compartments as well as 1378 and 1441 of B compartments were identified, respectively. Compared to the saline, a total of 14.74% of compartments were changed by the morphine, with 227 regions switching from compartment A to B and 164 regions switching from compartment B to A (Figure 2F). The A/B switching compartments were mainly attributed to several chromosomes, particularly for the Chr 2, 5, 8, 9, 12, and 18 (Figure 2G). These data revealed attenuated genome compartmentalization induced by morphine.

Since the change in compartmentalization closely correlates with the gene expression profile [23], we thus profiled genome-wide transcriptional expression in conjunction with A/B switching compartments. Among the significantly down-regulated 813 genes and up-regulated 664 genes (false discovery rate (FDR) < 0.05 and Fold change (FC) (morphine vs. saline) > 1.5) in the morphine group (Figure S4A), we discovered that the switching from compartment A to B caused 179 genes to be significantly dysregulated, and the switching from the B to A compartment caused 150 genes to be significantly dysregulated (Tables S6 and S7). The representative images of the A/B compartment switch and the corresponding significantly changed genes were visualized on Chr1 (Figure 2H). Next, all detected genes in the A/B switching compartments were further subjected to gene set enrichment analysis (GSEA) (with a reactome subset of canonical pathways). Different from the multiple enriched functional clusters using all significantly changed genes (Figure S4B), two gene sets named adaptive immune system (normalized enrichment score (NES) = −1.88) and mitogen-activated protein kinase (MAPK) family signaling cascades (NES = −1.69) were significantly enriched in the saline group at FDR < 0.25 (Figure 2I). Comparative analysis of enriched gene expression in morphine vs. saline based on the RNA-seq found that most genes in the adaptive immune system were down-regulated by morphine. Those switched compartmentalization-linked abnormal genes further demonstrated an impaired innate immune response following morphine administration [24]. Furthermore, the dysregulated genes involved in MAPK signaling reflected the development of morphine tolerance and dependence as previously reported [25]. Collectively, morphine appeared to attenuate genome compartmentalization, which may lead to the dysregulation of gene expression profiles of cortical neurons.

### Morphine modifies the size of TADs

TAD and TAD boundary are two basic features of the TAD organization [26]. To identify topological associating domains, we employed a Hidden Markov Model (HMM) on the directionality index (DI) from a Hi-C matrix to measure the level of upstream or downstream interaction bias for a genomic region [26]. By comparing the TADs across the entire genome of the two groups, we found that the number of TAD boundaries varied less than 4%, with 1222 and 1268 TAD boundaries called in saline and morphine groups, respectively (**Figure 3A**). Insulation score profiles of these two groups are highly correlated, with a Pearson correlation coefficient of 0.898 (**Figure 3B** and Figure S5A). Moreover, heatmaps presented a similar distribution of insulation scores around ±1 Mb TAD boundaries of these two groups (Figure 3C and Figure S5B). These data suggested that morphine exerted a minor impact on the insulation strength of TAD boundaries across the entire genome.

**Figure 3.**
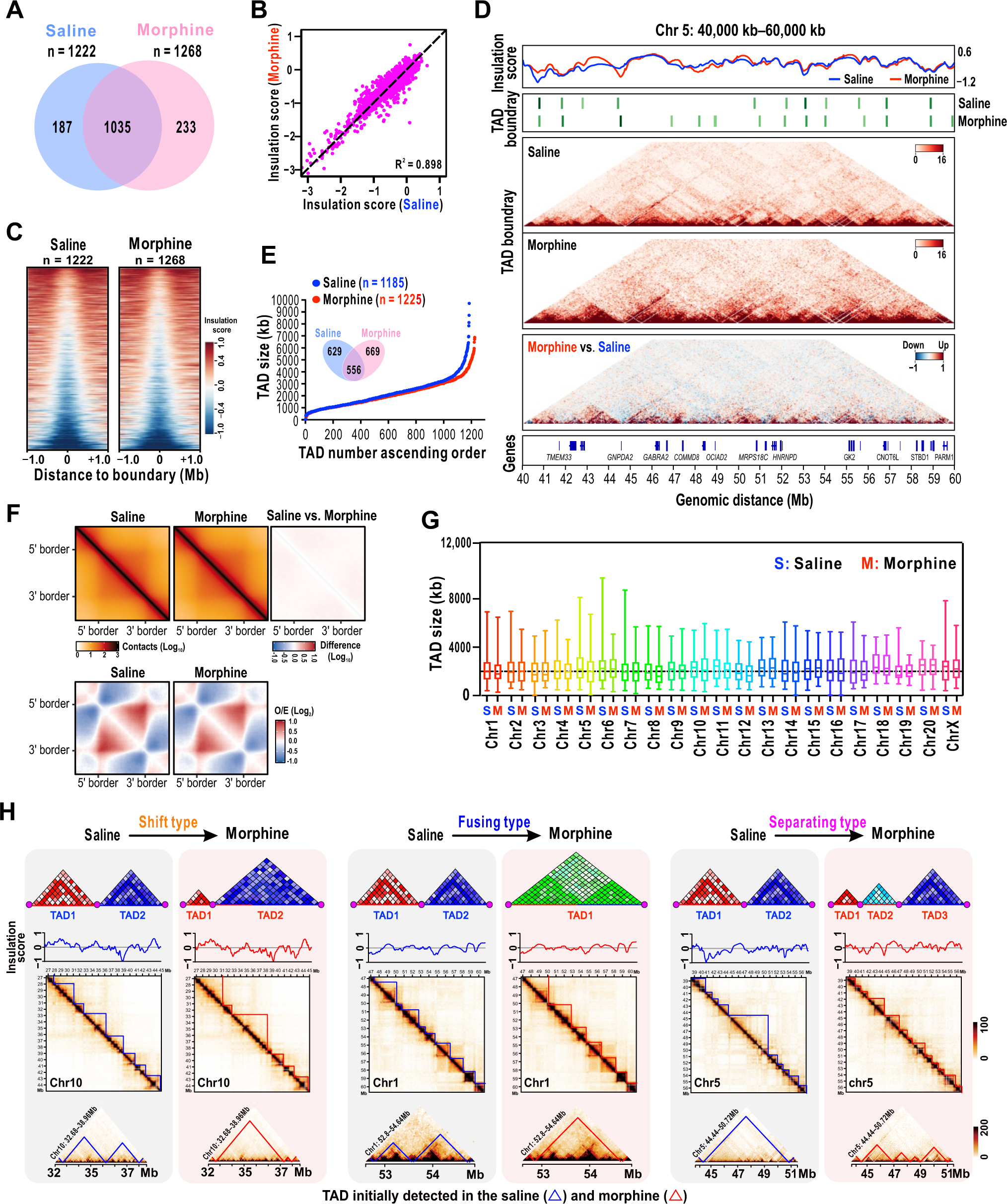
Morphine alters chromatin TADs. **A.** Venn diagram shows the number TAD boundary. **B.** Quantification of insulation score differences in saline versus morphine using the identified boundaries. **C.** Heatmaps of insulation scores centering around ±1 Mb TAD boundaries. **D.** An example of TAD and boundary alteration in response to morphine. A snapshot of insulation score curves in the saline (blue) and morphine groups (red), distribution of TAD boundary, contact maps, differential contact matrix, and distribution of genes (from top to bottom) was plotted for Chr5: 40 Mb–60 Mb. The differential contact matrices were generated by subtracting the normalized morphine matrix from the saline matrix. **E.** Scatter plots display the TAD size of all identified TADs across the entire. Inset: venn diagram shows the number. **F.** The aggregated at the center of the plot with ICE normalization (top) or distance normalization (bottom) via APA. G. Box–plot shows the size of identified TAD in each chromosome. **H.** Representative Hi-C contact maps display three altered TAD types in morphine. For each contact map, insulation score tracks are coupled. Domains emerging in the saline (blue) and morphine groups (red) are demarcated by color–coded lines. Bin size, 40 kb. The color bar denotes q-normed reads. Schematic diagrams of the different types of altered TADs caused by morphine are shown in the top lane. The circle with a magenta color represents the TAD boundary. The changed TADs appeared in the square are displayed in the restricted region. TADs, topologically associated domains; ICE, iterative correction and eigenvector decomposition; APA, aggregate peak analysis.

We next compared the location of identified TAD boundaries. If the location of an identified TAD boundary in the morphine group varied within two bins (80 kb) of the TAD boundary in the saline group, we defined it as the same TAD boundary; otherwise, we defined it as a different TAD boundary. Notably, the distribution of some TAD boundaries was clearly altered by morphine, including disappeared boundaries, neo-boundaries, and shifted boundaries (Figure 3D and Figure S5C). A total of 411 different boundaries were discovered by evaluating all recognized TAD boundaries across the entire genome, with 187 and 233 specific boundaries in the saline and morphine groups, respectively (Figure 3A).

The difference in TAD boundary motivated us to examine the effect of morphine on TAD. TAD boundaries in the saline and morphine groups separately insulated 1185 and 1225 TADs across the entire genome (Figure 3E and Figure S5D). The number of TADs was less affected not only across the entire genome but also at the individual chromosome (Figure S5E). The total TAD coverage across the entire genome was 39.22% in saline and 38.05% in morphine, ranging from 0.92% to 3.45% for individual chromosomes (Figure S5F). To gain further insight into whether topological architecture was influenced by morphine, we resized and aggregated all TADs genome-wide. The aggregated plot with iterative correction and eigenvector decomposition (ICE) normalization (top panel) showed no change, whereas the aggregated plot with distance normalization (bottom panel) presented slightly changed TADs in the morphine group (Figure 3F). By further comparing the size of TADs, we found that some TADs were altered by morphine (Figure 3E, G, and Table S8). For example, most TADs in Chr5 were clearly reduced in size, whereas some TADs in Chr10 were clearly enlarged in size (Figure S5G). As quantified in Figure S5H, morphine reduced the overall number of TADs with sizes above 4000 kb (> 4000 kb). In particular, six of those identified TADs with a size above 6900 Kb in the saline group were not observed in the morphine group (Figure 3E, Figure S6, and Table S8), suggesting that some TADs with large sizes might be reduced by morphine treatment. Moreover, the morphine group presented a smaller average size of TADs for the majority of chromosomes, with the exception of four chromosomes (Chr2, 6, 10, and 15) with the increased size of TADs and two chromosomes without alteration (Figure S5G and S6). These data suggested that morphine treatment might lead to the decreases in long-range genomic contacts. Collectively, morphine significantly altered both chromatin topological structure and the size of TADs at different degrees; however, morphine showed less impact on the insulation strength of the TAD border.

### Morphine causes different types of TAD alteration

Considering that morphine markedly altered the size of TADs, we continued to explore how these TADs were affected. Compared to the saline group, all TADs in the morphine group were classified into four types: unchanged, fusing, separating, and shift (Figure S7A). In the morphine group, over half of the TADs (54.65%) were altered. Among all those detected TADs in the morphine group, shift, separating, and fusing TADs accounted for 40.93%, 9.97%, and 3.76%, respectively (Figure S7A). In particular, both shift and separating TAD types in Chr5 took the most proportion of the altered TADs (Figure S7B). Among all the identified TADs in Chr5, 58.33% and 15.48% TADs were shifted and separated in the morphine group, respectively, with a small proportion (1.19%) of TADs fused (Figure S7A). In contrast to Chr5, the proportion of fusing type of TADs in Chr1 was much higher than that in other chromosomes (Figure S7B), whereas none of the fusing or separating TAD was found in Chr10. We proposed that morphine-extended TAD size in Chr10 attributed, to a large degree, to the shift of TADs (Figure S7A and B). The representative Hi-C contact maps marked with domains were presented to elucidate morphine-modified TADs (Figure 3H). Collectively, our data demonstrated that morphine remarkably impacts the chromatin TADs of cortical neurons.

### Morphine regulates the gene expression profile through altering TADs

To investigate how the altered TADs induced by morphine regulate gene expressions, we analyzed the transcriptional profile of genes around all identified TAD boundaries of cortical neurons. All genes from genome-wide RNA-seq were mapped to TAD boundaries, showing that the loci of 3160 genes were correlated to the TAD boundaries. Among them, the loci of 2663 genes were located in the same TADs of both groups; moreover, 312 and 185 genes were mapped to specific TADs in the morphine and saline groups, respectively (Figure S8A and Table S9). By contrast, 275 genes were significantly changed by morphine (FDR < 0.05 and FC > 1.5), including 186 down-regulated and 89 up-regulated genes (**Figure 4A**). The fragments per kilobase of exons per million mapped reads (FPKM) of 275 differentially expressed genes (DEGs) were clustered following the test conditions. Heatmap displayed the distinct expression patterns of these DEGs (Figure 4B).

**Figure 4.**
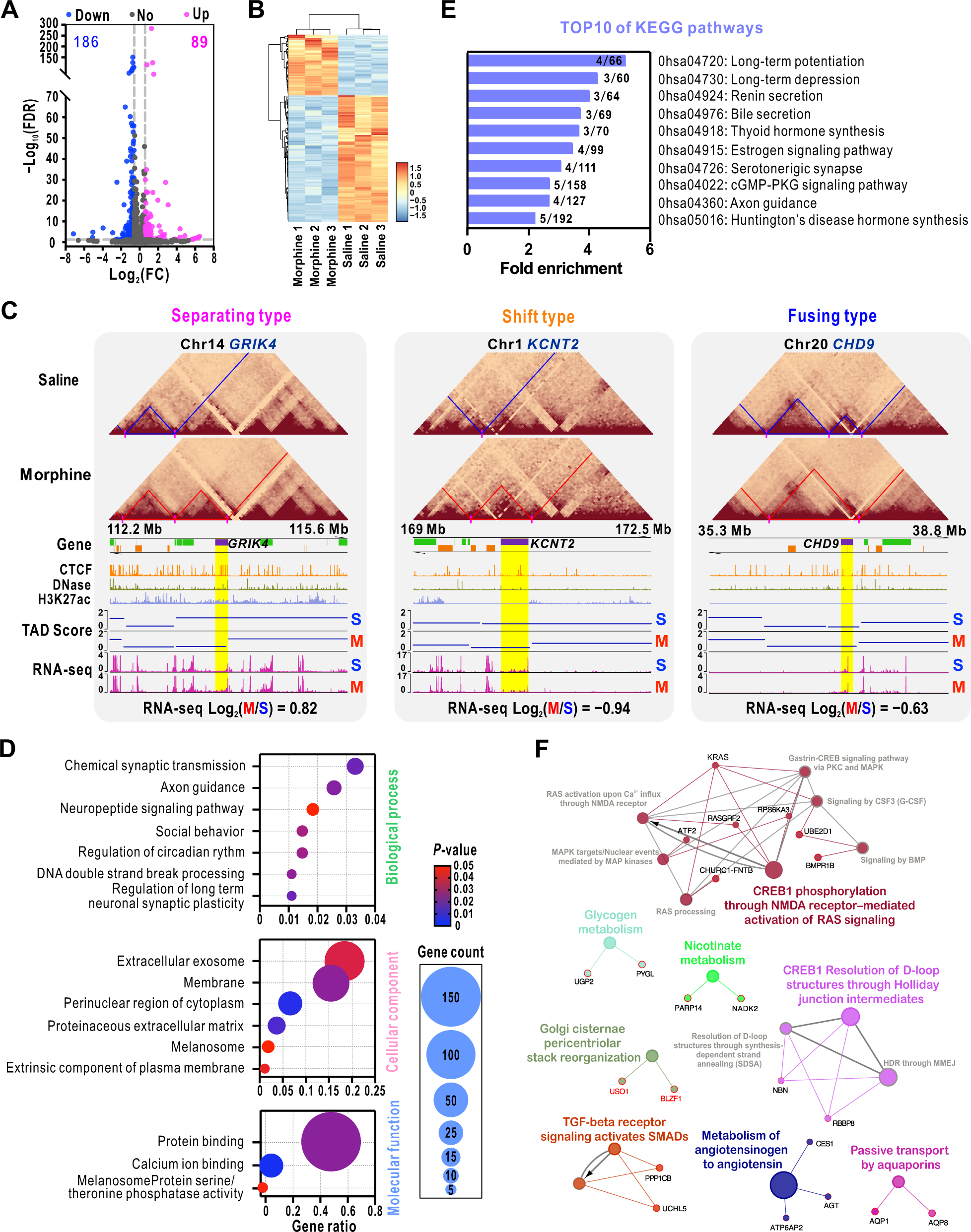
Morphine regulates the gene expression profile by altering TADs. **A.** Volcano plot shows the DEGs around the identified TAD boundaries. **B.** Heatmap of hierarchical clustering of all DEGs around TAD boundaries. Each column represents an experimental treatment, and each row represents a screened DEG. **C.** Representative DEGs around TAD boundaries to demonstrate the separating, shift, and fusing TADs. TADs emerging in saline (blue) and morphine (red) are demarcated by color-coded lines. Bin size, 40 kb. The rectangle shape with purple color represents DEG around the altered TAD boundary. IGV screenshots of CTCF, DNase, and H3K27ac ChIP-seq peaks are presented with different colors. **D.** The bubble chart shows the significantly changed terms classified in three aspects of GO enrichment analysis. **E.** The bar chart shows the top 10 most-enriched KEGG pathways of DEGs in response to morphine. **F.** Network enrichment analysis of DEGs. The color code of nodes corresponds to the functional group to which they belong. DEGs, differentially expressed genes; IGV, integrative genomics viewer; ChIP-seq, chromatin immunoprecipitation-sequencing; CTCF, CCCTC-binding factor; GO, gene ontology.

CCCTC-binding factor (CTCF) and multiple transcription factors (TFs) are enriched at the nearby TAD boundaries, and play critical roles in the correct insulation of two neighboring TADs and gene expression regulation [27]. To test whether identified TAD boundaries were enriched with the chromatin immunoprecipitation-sequencing (ChIP-seq) peaks, we mapped our list of TAD boundaries to publicly available CTCF, DNase, and H3K27ac ChIP-seq datasets of rhesus macaque (GSE163177 and GSE67978) [28,29]. The results presented that the identified TAD boundaries significantly enriched ChIP-seq signals (Figure S8B), indicating the relevance of those altered TAD boundaries in response to morphine treatment to the dysregulated gene expression profile. Among the 275 DEGs, a total of 95 and 54 genes were located in specific boundaries in the morphine and saline groups, respectively (Figure S8C). The representative up- or down-regulated genes around the specific TAD boundaries as exemplified by the changes were visualized in Figure 4C. The boundary formation around *GRIK4* loci led to a separation of the original TAD in the saline group into two TADs in the morphine group. We hypothesized that in response to long-term morphine treatment, the small neo-TAD altered chromatin topological architecture of cortical neurons, which may promote the transcriptional activation of *GRIK4* genes. In addition, the shift of the TAD boundary around *KCNT2* loci altered the transcriptional activity of genes, which could explain the morphine-induced downregulation of *KCNT2*. Furthermore, two detected TADs in the saline group were fused to one TAD in the morphine group. This fusing around *CHD9* loci mediated the conversion of the transcriptionally activated *CHD9* gene to an inactivated state (Figure 4C and Figure S8D). Taken together, morphine promoted the formation of new TADs, the disappearance or shift of original TADs, and then probably altered the interaction of regulatory elements with their cognate genes, thus modifying the transcriptional profile of genes.

All DEGs around TAD boundaries were further subjected to gene ontology (GO) enrichment analysis using DAVID v6.8 online server. For biological processes, several neuron-associated processes were significantly enriched, including synaptic transmission, neuroplasticity, and axon guidance. For cellular components, a membrane-bound pattern was profoundly enriched, such as extracellular exosome (Figure 4D). Intriguingly, the top 10 pathways from KEGG pathway enrichment revealed the association of DEGs with long-term potentiation (LTP) (0hsa04720), long-term depression (LTD) (0hsa04730), cGMP-PKG signaling (0hsa04022), and axon guidance (0hsa04360) (Figure 4E), which have previously been demonstrated to play important roles in synaptic plasticity [30,31].

All DEGs around TAD boundaries implicated in cell signaling were visualized via ClueGO/CluePedia plugin from Software Cytoscape (version 3.8.2). Importantly, several signaling pathways were found to be modified by morphine, including the N-methyl-D-aspartate (NMDA) receptor-mediated cAMP response element-binding protein 1 (CREB1) phosphorylation, pathway network for CREB1 resolution of D-loop structures through Holliday junction intermediates, and the transforming growth factor beta (TGF-β) receptor signaling (Figure 4F). Indeed, the critical role of CREB phosphorylation via NMDA receptor has been highlighted during the process of diverse drug addiction [32]. We believed that altered TADs may be critical for modulating the gene expression profiles involved in synaptic plasticity, which may contribute to morphine effects such as tolerance and dependence.

### Morphine modifies chromatin looping

The formation of loops brings pairs of genomic regions that lie far apart along the linear genome together in the space [33]. We identified DNA looping using the major Hi-C loop-calling tool HiCCUPS from Juicer according to the previous reports [34]. The results from the quantitative analysis showed that 5507 and 8425 DNA loops were called in the saline and morphine groups, respectively (**Figure 5A**). Except for Chr5 with a reduced number of loops, the other chromosomes in the morphine group consistently showed an increased number of loops (Figure 5B and Figure S9A). A similar trend was observed in the number of differential DNA looping events (Figure 5B and Figure S9B). The exemplified coverage-corrected Hi-C contact matrices separately presented an increased DNA looping event (black dots) of Chr1 and a decreased DNA looping event (black dots) of Chr5 in the selected regions (Figure 5C). The quality of all identified DNA loops was assessed by aggregate peak analysis (APA). Both APA plots presented the intense center pixels surrounded by less intense pixels, and the score of APA plots was quite high, with 2.62 in the saline group and 2.99 in the morphine group, respectively (Figure 5D), indicating the accuracy of the identified DNA loops.

**Figure 5.**
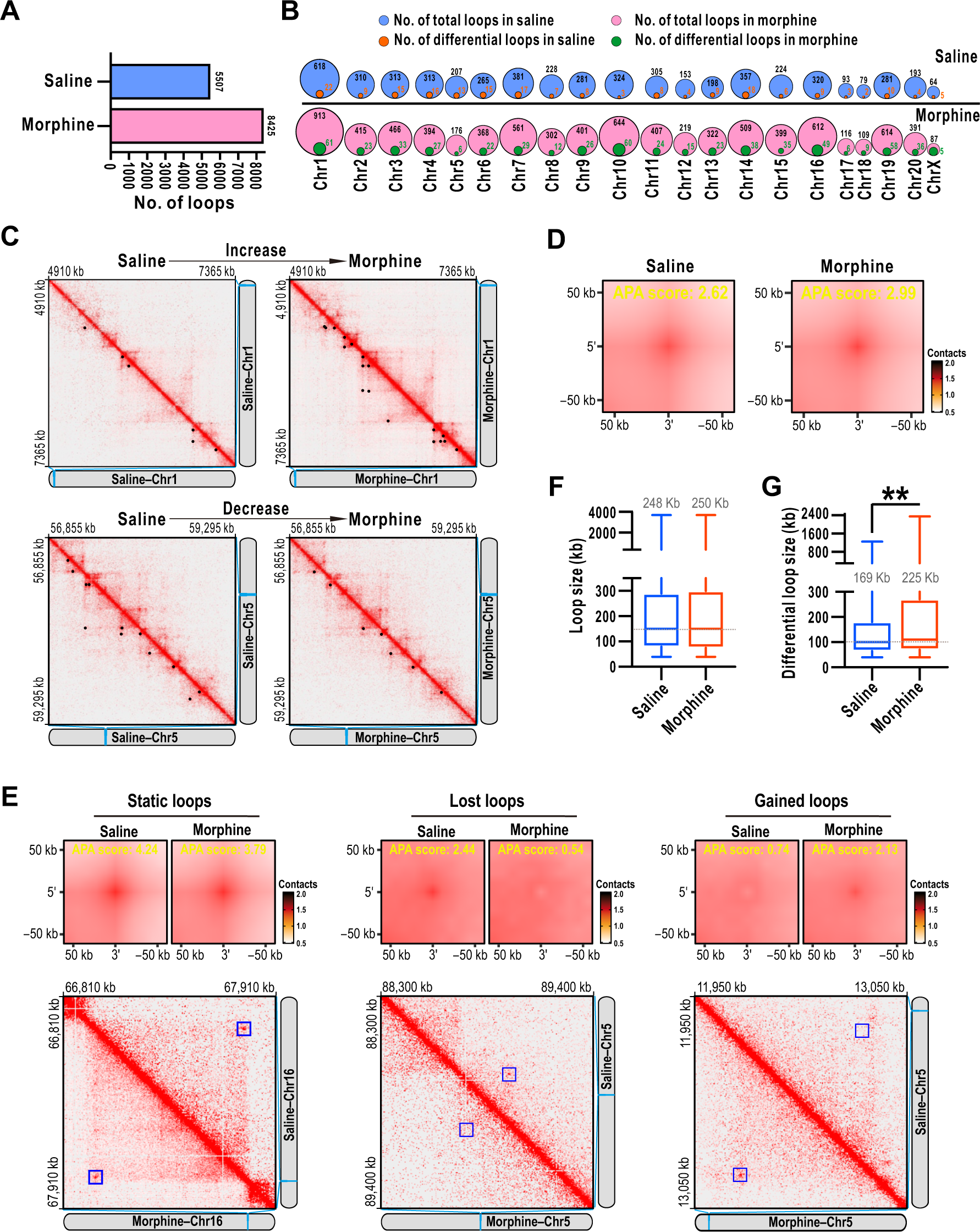
Morphine regulates chromatin looping events. **A.** Bar graph of identified loops across the entire genome in the saline and morphine group. **B.** Venn diagrams show the number of chromatin loops and differential loops in each chromosome. The size of the circle represents the number of loops, and values inside or around the circle are the number of identified loops. **C.** Representative coverage–corrected Hi-C contact matrices show the increased and decreased chromatin loops in morphine. Loops are marked by black spots. The matrices were plotted using Juicebox. **D.** APA plots show the aggregated signal across all identified chromatin loops. The score of aggregated signals was displayed in yellow color. **E.** APA plots for morphine-induced static, lost, and gained loops, which are marked by a blue square. **F.** Box-plot shows the size of chromatin loops across the entire genome. The average loop size is shown on the top of the box. **G.** Box–plot shows the size of differential loops. The average loop size is presented in the middle of the box. Student’s t-test, \*\**P* < 0.01.

Compared to the saline group, all detected DNA loops in the morphine group were classified into three categories: static loops (loops identified in both groups), lost loops (loops only detected in the saline but not the morphine group), and gained loops (loops only detected in the morphine but not the saline group). APA plots of these types of DNA loops showed a clear difference in contact frequencies between the two groups, indicating that the identified differential loops were entirely lost or gained by morphine. Moreover, the representative matrices presented the static, lost, and gained DNA loops, respectively (Figure 5E). These findings further indicated that morphine caused a remarkable re-arrangement of chromosome conformation, which was compatible with the alteration in TADs.

Close analysis of loop length showed no apparent difference in the average size of all loops (248 Kb in the saline group and 250 Kb in the morphine group) (Figure 5F). We then compared the loop size of all differentially gained and lost loops. Interestingly, the average size of 203 differential loops in the saline group was 169 Kb, whereas the average size of 597 differential loops in the morphine group was dramatically enlarged to 225 Kb (Figure 5G, Tables S10, and S11). These results indicated that morphine promoted not only the formation of neo-loops but also the extension of differential loops, implying enhanced long-range contacts of regulatory elements with associated target genes.

### Altered DNA loops modulate the expression of target genes associated with morphine effects

We wondered which genes were modulated by the altered DNA loops induced by morphine. All up- and down-regulated genes were separately mapped to the differential loops of individual chromosomes (**Figure 6A**). The results showed that 85 genes were significantly dysregulated (Padj < 0.05 and FC > 1.5) by morphine, with 33 genes upregulated and 52 genes downregulated (Figure 6B). Among these DEGs, 27 genes were attributed to the saline group and 58 genes were attributed to morphine treatment. Heatmap showed a distinct expression pattern of DEGs, illustrating a perfect cluster between the two groups (Figure S10A, Tables S12, and S13). The significant clustering of CTCF, DNase, and H3K27ac ChIP-seq peaks near the identified loop anchors further implied the correlation of those differential loops to the dysregulated genes (Figure S10B). We then chose 25 genes from DEGs that were mapped to differential loops to validate their changes in mRNA levels using an RT-qPCR assay. Importantly, the mRNA levels of most of these genes matched the results of the RNA-seq analysis (Figure 6E and Figure S10C), demonstrating that altered chromatin looping indeed alters the expression of the target genes after morphine treatment. Then, GO analysis using all DEGs found that two functional clusters, protein-protein/nucleotide binding, and immunity, were enriched. By analyzing the genes involved in each GO term (Figure 6C), we found that immunity-associated genes, *SYK*, *TK2*, *CSF1R*, *STK36*, and *MYH3*, were markedly up-regulated by morphine, whereas protein-protein/nucleotide binding-associated genes, *ZAP70*, *ABCA1*, *ATP4A*, *ABCG2*, and *FGR*, were significantly down-regulated. Intriguingly, a few DEGs in the morphine group were disclosed to be involved in several signaling pathways, such as epigenetic regulation of gene expression, neurotransmitter release cycle, anti-inflammatory response, opioid signaling, MAPK signaling, and cellular senescence (Figure S10D). In the saline group, 27 DEGs were not linked to the associated cell signaling, but GO ontology analysis of these DEGs discovered multiple biological processes dysregulated by morphine, such as protein phosphorylation and cellular response to extracellular stimulus (Figure S10E).

**Figure 6.**
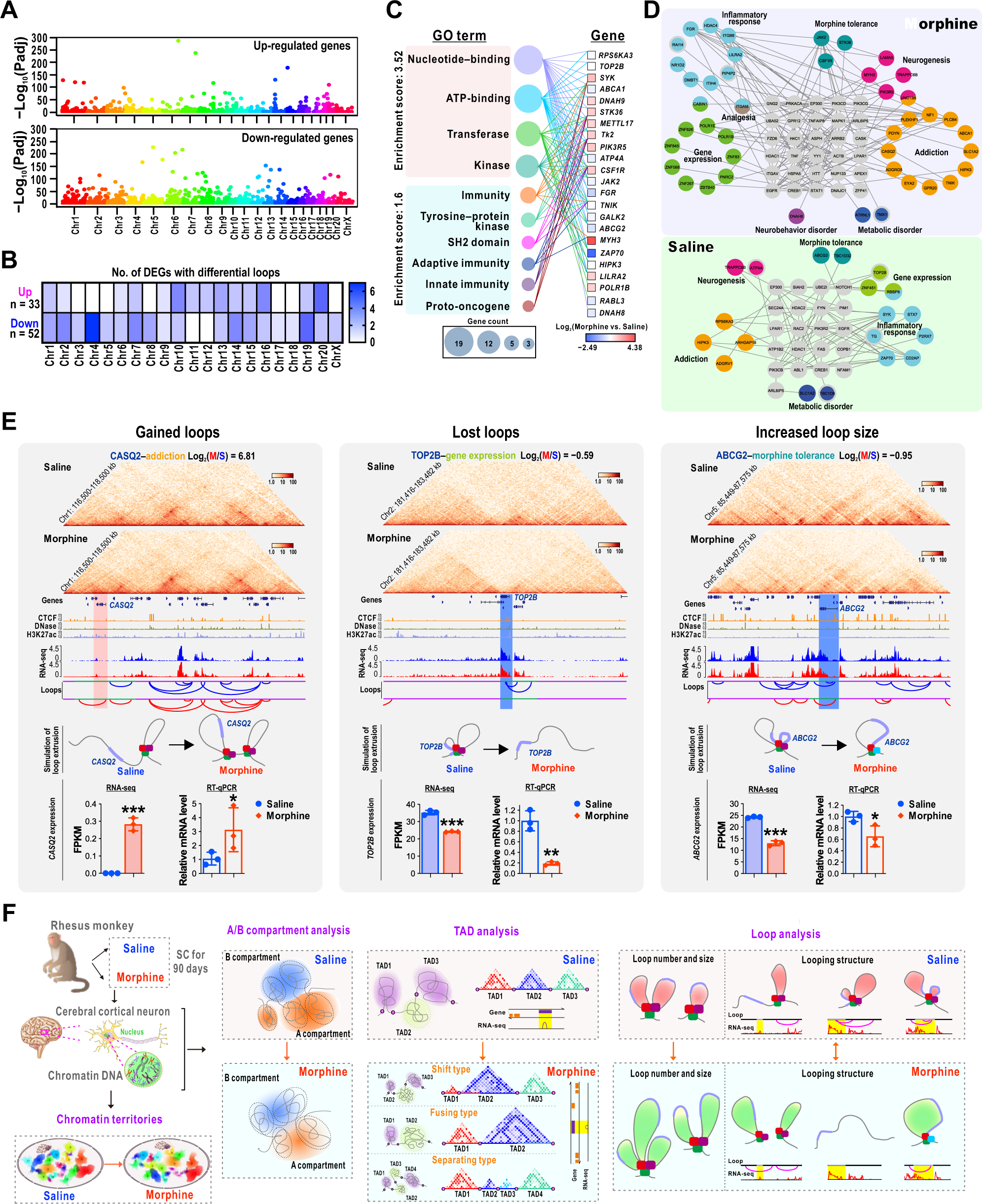
Morphine-induced alteration of DNA loops dysregulates the target genes associated with the morphine effects. **A.** Manhattan plots show the significantly up- and down-regulated genes in each chromosome. Each point represents a single gene, with physical position (chromosome localization) plotted on the x-axis and −Log_10_ (Padj) on the y-axis. **B.** Heatmap presents the number of DEGs linking to differential loops in each chromosome. **C.** Function annotation of DEGs in the top two GO terms. The size of the bubble represents the number of enriched genes. **D.** PPI network of annotated DEGs. Each functional category is color–coded, and the interactions between two proteins are linked with a gray line. **E.** Representative genes are modified by the change of chromatin looping architecture induced by morphine. Top two rows: the Hi-C contact maps were rotated 45^∘^ so that the main diagonal is horizontal; IGV screenshots of CTCF, DNase, and H3K27ac ChIP-seq peaks are presented with different colors. The location of DEG linking to the differential loop at linear genome is marked with light purple in simulated loop extrusion. The region for the simulation of loop extrusion is covered with a green line at the x-axis of loop calls. For the bar graphs of RNA-seq and RT-qPCR in the last row, data from three replicates are presented as mean ± SD. Student’s t-test, \**P* < 0.05, \*\**P* < 0.01, \*\*\**P* < 0.001. **F.** The simplified schematic model shows the impact of long-term morphine on chromatin architecture. PPI, protein–protein interaction network; RT-qPCR, real-time quantitative PCR; FPKM, the fragments per kilobase of exons per million mapped reads.

Lastly, the corresponding proteins on the DEGs were annotated, and the protein–protein interaction network (PPI) was investigated through Reactome [35]. PPI analysis uncovered that over 70% of annotated proteins (43/58 in the morphine group, 19/27 in the saline group) were involved in several functional categories, including drug addiction, morphine tolerance, analgesia, neurobehavior, inflammatory response, gene expression, metabolic disorder, and neurogenesis (Figure 6D). Interestingly, a few proteins, such as CREB1 and histone deacetylase 1 (HADC1) which are critical players in drug addiction, were uncovered to interact with those differentially annotated proteins [32]. These data indicated that the altered conformation of the DNA loop played a critical role in regulating the expression of genes involved in morphine effects, such as addiction, tolerance, and neurobehavior.

### Morphine causes altered DNA looping events that regulate the transcription of target genes

To clarify how the changes in differential loops regulated gene expression profiles, we investigated all looping events around the significantly changed gene loci. Compared to the saline group, the formation of a neo-DNA loop (gained loop) at the *CASQ2* loci greatly activated the transcriptional expression of *CASQ2* in the morphine group (Figure 6E). In addition, those genes with large sizes at the linear genome maps were not expressed even though part of their sequence was situated in a looping architecture. For instance, the length of the *SNX29* gene is 589 kb at the linear dimension, and three loops spanning the SNX29 sequence detected in the saline group were markedly altered by morphine, resulting in the formation of another neo-loop between the original loops. As a result, this alteration of DNA spatial architecture promoted the transcription of *SNX29* gene (Figure S11). In the opposite condition, the looping structure around *TOP2B* loci in the saline group disappeared in response to morphine. The absence of this DNA spatial architecture reduced the contacts of regulatory elements, ultimately inhibiting the transcription of *TOP2B* (Figure 6E). Apart from such huge changes in loop structure, micro-changes in looping structures, such as an alteration in looping architecture at *TMEM114* loci, also modulated gene expression. We suppose that altered DNA looping structure induced by morphine may modify the activities of relevant transcriptional regulatory elements, thus up-regulating TMEM114 transcription (Figure S11).

Morphine not only altered the number of looping events but also caused an extension of looping architecture, indicating the occurrence of long-range regulation (Figure 5). For example, morphine caused an extension of the looping structure around *ABCG2* loci, resulting in a decreased transcription of the *ABCG2* gene (Figure 6E). Similar to the looping structure around *ABCG2* loci, an original small loop around the *ZAP70* loci in the saline group was mapped to a large loop structure in the morphine group, causing a decreased transcriptional activity of the *ZAP70* gene (Figure S11). Collectively, different types of alteration in chromatin looping architecture caused by morphine can differentially regulate the gene transcription activities of cortical neurons.

## Discussion

Chronic morphine administration has far-ranging consequences beyond analgesia and dependence. The aberrant gene expression at transcriptional, translational, and epigenetic levels induced by morphine has been studied in diverse animal models [1,2,36]. However, how morphine regulates the transcriptional activity of target genes at the level of chromatin 3D architecture is unknown. Here, by combining Hi-C technology and genome-wide transcriptional analysis, we revealed a disorganized chromatin architecture in the multi-hierarchical structure of cortical neurons in the rhesus monkey with high genetic similarities to the human genome [37]. On the macro scale, morphine re-arranged chromatin territories in the nucleus of cortical neurons. At higher resolution, the genome-wide chromatin compartmentalization was slightly attenuated, with a total of 391 switching compartments. Over half of the TADs across the entire genome were modified, including shift, separating, and fusing. Notably, morphine promoted not only the occurrence of looping events but also long-distance interaction (Figure 6F). Those DEGs associated with altered chromatin architecture were mainly enriched in the several signaling pathways related to neuroplasticity, synaptic receptor transmission, and inflammation. Our findings provide a pivotal clue connecting an altered chromatin 3D architecture, a regulatory mode of gene expression as well as morphine effects.

Individual chromosomes preferentially occupy separate territories which are associated with both intra- and inter-chromosomal compartments [38]. Within each compartment, TADs constrain chromatin interactions. Within each TAD, loop extrusion may make it easier for region-specific enhancer-promoter interactions, protecting against the overall transcription environment [39,40]. The competition between compartmental phase separation and nonequilibrium active loop extrusion leads to the emergence of chromatin organization on the megabase scale [41–43]. Chromatin compartmentalization is mainly based on the active and inactive states of local chromatin, and the same compartments tend to close together in space [41]. Hence, the A compartment is frequently positioned in the euchromatin regions and interior nuclear space, whereas the B compartment is largely located with heterochromatin regions and nuclear lamina-associated domains. In this study, we found that long-term morphine administration caused an increase in looping but weakened genome-wide compartmentalization. The attenuated compartmentalization induced re-arrangement of the chromosome territories, which was mostly associated with increased intra-chromosomal interaction. For example, the enhanced intra-chromosomal interaction of chr1 caused attenuated compartmentalization but facilitated loop extrusion. Although both intra-chromosomal interaction and loop extrusion of Chr5 were reduced by morphine, the inter-chromosomal interaction of Chr5 was mostly promoted, suggesting an increased contact probability of Chr5 with the other chromosomes [44].

Growing studies have demonstrated that the disruptions or disorganization of TAD boundary can cause large-scale structural variations, contributing to abnormal gene expressions and eventually a molecular pathological mechanism of human disease [45–48]. For instance, enhancer adoption results from the depletion of a TAD boundary at the *LMNB1* loci, promoting *LMNB1* transcription and eventually leading to the progress of neurological disorder [49]. In this study, despite less impact on the number of TAD boundaries and boundary strength, morphine altered around one-third of TAD boundaries, with 18% specific TAD boundaries. As both insulation strength and spatial distribution are two important features of TAD boundary [50], we considered that altered TAD boundaries may contribute to the dysregulated gene expression. Through mapping identified genes to the identified TAD borders, we discovered that multiple target genes located around specific TAD boundaries were dysregulated by morphine. For instance, a small TAD was detected around *GRIK4* loci, which may accelerate the interaction of regulatory elements with *GRIK4* and thus promote its transcription. *GRIK4* is a gene encoding a high-affinity kainate receptor (KAR) subunit, GluK4. Gain of function of this gene induces severe depression, anxiety, and reduced locomotor activity in GluK4^over^ mice [51]. *GRIK4* variants have also been discovered in patients with acute postoperative pain and excessive morphine use [52]. Our findings indicate that altered chromatin 3D structure may contribute to the regulation of gene expression in the cortical neurons exposed to morphine. Through GO analysis of specific TAD boundaries associated DEGs, we found that a few signaling processes associated with neuronal activities, structural plasticity, and neurobehaviors were dysregulated by morphine.

The critical role of CREB-mediated signaling pathways in neurons has been demonstrated in various intracellular processes, such as long-term synaptic potentiation, neuronal plasticity, and drug addiction [32,53]. Phosphorylation-activated CREB mediates the transcription of target genes, such as brain-derived neurotrophic factor (BDNF), and ultimately forms the reward memories for abused drugs [54]. Importantly, our data discovered that several DEGs induced by TAD alteration modulate CREB activation through NMDA receptors after chronic morphine treatment. Indeed, morphine exhibits a bidirectional impact on gamma-aminobutyric acidergic (GABAergic) synaptic plasticity, including inhibiting presynaptic LTP and preventing LTD [55,56]. Consistently, by KEGG pathway enrichment analyses, we discovered that some of the TAD alteration-associated genes are involved in synaptic LTP and LTD, supporting a notion that morphine-induced bidirectional GABAergic plasticity reflects the neural adaption necessary for addictive properties of opiates [57]. In addition, the enriched cGMP-PKG signaling pathway has also been demonstrated to participate in the development of morphine tolerance. Inactivation of this pathway is predicted to be a promising strategy to avoid morphine tolerance during the treatment of neuropathic pain [58]. Taken together, morphine causes a genome-scale of topological structural alteration, and these changes in fundamental regulatory units alter the contact of regulatory elements with its locally targeted genes, thus regulating the transcription of these genes. This mechanism based on topological chromatin domains is of great significance to elucidate the complicated pharmacological and toxicological effects of morphine, such as analgesic effect [59], tolerance [60], inflammatory response [61], and addiction [62].

Less than 2% of human genome is thought to encode functional proteins, and the rest over 98% of human genome sequences regulate genes hundreds of thousands of base pairs away via forming DNA loops [63]. Loop-based transcriptional regulation is dynamically varied along with biological contexts. Indeed, our results also showed that morphine promoted the formation of DNA loops, leading to a dramatic boost in loop number; moreover, some loops were neo-formed while some loops were lost. More interestingly, quantitative analysis of these differential loops revealed that morphine markedly enlarged the size of the differential looping structure. These results indicate that morphine promotes not only the formation of loops but also the long-range interaction of regulatory elements with the target genes, revealing a novel mechanism by which morphine regulates the chromatin looping events.

There are 3-fold more differential loops in the morphine group versus the saline group in this study. In eukaryotes, there are primarily three different kinds of chromatin loops. Depending on their function, these loops are formed and maintained by different mechanisms: 1) loops that help the chromatin pack into mitotic or meiotic chromosomes to ensure accurate genetic information distribution; 2) loops that keep the genome functional and ensure precisely gene regulation; and 3) loops generated by continuous, intense transcription [64]. We thus proposed that not all differential loops in the morphine group were involved in the transcription of genes. Some chromatin loops are reported to be involved in temporarily or permanently suppressed genes. Hence, loops may be generated to either boost or inhibit the gene expression [65]. Even for those gene regulation-involved loops, not all genes were actively expressed. Therefore, our finding presented that ∼61% of DEGs were downregulated in response to morphine, suggesting that the formation of some looping might not correlate positively with gene activity. Lastly, we also analyzed the proportion of enhancer-gene loops based on the H3K27ac ChIP-seq peaks, and the results showed that the proportion of enhancer-gene loops in morphine-specific loops was higher than that in the saline and morphine group (data not shown). Additionally, we further mapped all up-regulated DEGs to the enhancer-gene loops. Consistent with the increased proportion of enhancer–gene loops, the proportion of up-regulated DEGs was indeed higher in the morphine-specific enhancer-gene loops than that in the enhancer-gene loops of saline, indicating that the altered enhancer–gene loop extrusion following chronic morphine administration indeed promoted gene expression.

The functional cluster analyses of the DEGs mapping to differential chromatin loops found that the most enriched functional clusters were related to protein binding. In the protein-binding functional cluster, *TOP2B* [66]*, ZAP70* [67], *ABCA1* [68], *ABCG2* [69], and *DNAH8* [70] have been shown to participate in the neurogenesis, inflammatory response, drug addiction, morphine tolerance, and/or neurobehavioral disorder. Although less evidence shows the direct roles of DEGs in the morphine-regulated process, some DEGs encoding proteins, such as G protein-coupled receptor 20 (GPR20) [1] and the G-protein effector neurofibromin 1 (NF1) [71], have been shown to participate in morphine dependence. Intriguingly, some DEGs encoding proteins, such as prodynorphin, are involved in the dependence of other addictive drugs [72]. It is worth noting that the looping architecture of some DEGs with unknown roles was discovered to be altered by morphine for the first time. For example, an activated transcriptional activity of a recently reported new gene *SNX29* is supposed to be caused by the formation of an extended loop between the original discontinuous loops, hinting its involvement in the morphine effect [73]. Further experiments will be required to elucidate this point.

Collectively, by investigating genome-wide chromatin architecture of non-human primate cortical neurons, a series of known or unknown morphine-regulated genes are proved to be altered along with DNA 3D architecture. Our finding provides critical hubs connecting chromosome spatial structure and gene networks associated with the morphine effect.

## Materials and methods

### Animals

Four male rhesus monkeys (*Macaca mulatta*; 3–4 years old; weighing from 3.0 to 5.0 kg) were purchased from Sichuan Green-House Biotech Co., Ltd (China). All monkeys were born by different mothers, but each two of them came from the same father to reduce the differences in genetic background. All monkeys were individually housed in stainless cages locating the same room under controlled conditions of humidity (40%–70%), temperature (23 ± 3°C), and light (12 h–light/12 h–dark cycle). All monkeys were fed with commercial monkey biscuits twice a day with free access to water. Moreover, fresh fruits and vegetables were provided once a day. The bodyweight of all monkeys was recorded once a week just before each feeding in the morning.

### Drug

Morphine hydrochloride was obtained from Northeast Pharmaceutical Group Co., (Cat#3557/12/25, China). Morphine was dissolved in 0.9% saline (sodium chloride) with a final concentration of 10 mg/ml.

### Experimental procedure

To reduce genetic differences, we selected four male monkeys from two fathers, and each pair of male monkeys from the same father was divided into saline and morphine (n = 2/group). Morphine was injected subcutaneously (SC) into the back legs of the monkey three times daily (at 9:00, 14:00, and 21:00) for 90 days continuously to produce dependence. The doses of morphine were gradually elevated as following paradigm: day 1–7: 3 mg/kg, day 8–14: 6 mg/kg, day 15–21: 9 mg/kg, day 22–28: 12 mg/kg, day 29–90: 15 mg/kg [2]. Monkeys in the saline group were injected SC three times daily with the same volume of 0.9% saline (0.5 ml/kg) as a control.

The criteria for physical dependence development of morphine is abrupt or spontaneous withdrawal, which was assessed as previously described [74]. Fourteen hours after the final morphine or saline injection, the precipitated withdrawal signs of all monkeys were scored once during each of five consecutive 30 min observation periods. The withdrawal signs evaluated included the followings: lying on the side or abdomen, drowsiness (sitting with eyes closed and lethargic or being indifferent to surroundings), fighting, avoiding contact, vocalizing, crawling and rolling, restlessness (pacing), ptosis, tremors, retching, vomiting, coughing, vocalizing when abdomen palpated, rigid abdomen, salivation. The observer was “blind” regarding the assignment of treatments. Differences between saline and morphine groups were measured by GraphPad Prism 9 software (version 9.5.0) using two-way ANOVA. The *P* < 0.05 was considered statistically significant.

### Preparation of cerebral cortical cells

Twenty-four hours after the final dosing, monkeys were anesthetized with pentobarbital sodium and the brain was removed. Cerebral cortical gray matter was carefully dissected away from the white matter on the ice and immediately dissociated using an Adult Brain Dissociation Kit (Cat#130-107-677, Miltenyi Biotech) according to the manufacturer’s instruction. In brief, the dissected fresh cerebral cortex was cut into small pieces with a scalpel in cold dulbecco’s phosphate buffered saline (D-PBS). After centrifugation, the pellet was harvested, and an appropriate volume of enzyme-supplemented digestion solution was added. The resuspended mixture was aspirated into gentleMACS C tubes (Cat#130-093-237, Miltenyi Biotech) and then the cerebral cortical pieces were mechanically dissociated with the gentleMACSTM Octo Dissociator with Heaters (Cat#130-096-427, Miltenyi Biotech). The dissociated mixture was filtered with a MACS SmartStrainer (70 μm) (Cat#130-098-462, Miltenyi Biotech) to remove cell clumps or cells with a diameter > 70 μm. Lastly, myelin and cell debris in the dissociated cells were removed using the Debris Removal Solution (Cat#130-109-398, Miltenyi Biotech), and red blood cells were lysed with Red Blood Cell Lysis Solution (10×) (Cat#130-094-183, Miltenyi Biotech) as the manufacturer’s instructions.

### Isolation of cortical neurons by flow cytometry

The enrichment of neuron cells from the fresh cortex of rhesus monkey was referred to the previously reported methods with less revision [75]. Cells were orderly fixed and permeabilized with Fixation Buffer and Permeabilization Buffer coming from the Foxp3/transcription factor staining buffer set (Cat#00-5523, Thermo Fisher Scientific) according to the manufacturer’s instruction. Then, the permeabilized cells were immunostained with the Alexa Fluor 488 conjugated anti-NeuN antibody (1:100; Cat#MAB377X, Millipore) at 4 °C for 30 min in the dark. After washing, the cell pellet was collected and further stained with DAPI (1:10,000; Cat#C0060, Solarbio) at 4 °C for 5 min in the dark. Lastly, the neurons were identified and sorted by FACSAria SORP (BD Biosciences) with appropriate gating parameters, and data analysis was performed using FlowJo 10 software (Tree Star, San Francisco, CA).

### Preparation of DLO-HiC sample

The preparation of the DLO Hi-C sample and the related data analysis mainly referred to the previously reported procedure [19].

#### Restriction enzyme digestion

The sorted neurons were pelleted from gray matter tissue after the purification step and washed once with pre-cold phosphate-buffered saline (PBS). Neurons were resuspended with pre-cold PBS. Next, 37% formaldehyde (Cat#252549, Sigma) was directly added into re-suspended cells with a final concentration of 1% and precisely cross-linked at room temperature for 10 min to cross-link all chromatin DNA. After cross-linking, the excess formaldehyde was quenched with glycine by incubating for 5 min at room temperature. All neurons were collected via centrifugation at 2000 r.p.m. for 5min. The cross-linked neurons were lysed in lysis buffer (10 mM NaCl, 10 mM Tris-HCl pH 8.0, 0.2% sodium dodecyl sulfate (SDS), 0.3% Igepal CA-630, and protease inhibitor (Roche)), and lysed at 50 °C for 5 min. Then, all nuclei were pelleted by centrifugation at 1000 r.p.m. for 5 min and washed once with ice-cold PBS. Lastly, the cross-linked chromatin was digested with *MseI* (Cat#R0525L, NEB) in NEB buffer for 6 h at 37 °C with simultaneous rotation at 15 r.p.m..

#### MmeI half-linker ligation

After digested with a restriction enzyme, 50 μl of T4 ligation reaction mixture containing 300 ng/μl half-linkers (Linker 1: 5’-p-TAGTCGGAGAACCAGTAG-3’, Linker 2: 5’-CTAGCTACTGGTTCTCCGAC-3’), 50 mM ATP, 2.5 units/μl T4 DNA ligase (Cat#15224025, Thermo Fisher Scientific) was added to 400 μl digested chromatin with thorough mixing. The reaction mixture was subsequently incubated at 25 °C for 1 h with simultaneous rotation at 15 r.p.m. Then, the nuclei pellet was harvested via centrifuging at 500 r.p.m. for 5 min. Lastly, the nuclei were washed twice with cold PBS.

#### In situ proximity ligation

The linker1-ligated and linker2-ligated nuclei were firstly resuspended in T4 DNA ligation solution (Cat#B69, Thermo Fisher Scientific) containing 0.5 units/μl T4 polynucleotide kinase (Cat#0201L, NEB). The reaction mixture was incubated at 37 °C for 30 min to phosphorylate all fragmented ends. Then, a T4 DNA ligation buffer containing 0.5 units/μl T4 DNA ligase (Cat#15224025, Thermo Fisher Scientific) was added to the reaction mixture. Lastly, ligate linker 1 or linker 2 containing fragments was incubated at 20 °C for 2 h with simultaneous rotation at 15 r.p.m..

#### Reversal of cross-linking and DNA purification

After in situ ligation, pellet nuclei were centrifuged at 1000 r.p.m. for 5 min, and the nuclei pellet was resuspended with ddH_2_O. A proteinase K digestion mixture was added, with a final of 0.5 mg/ml proteinase K (Cat#908239450-01-6, Sigma), 34.67 mM SDS, and 250 mM NaCl. After incubation at 65 °C for 2 h, an equal volume of phenol: chloroform: isoamyl alcohol (25: 24: 1) was added to the sample with vigorously shaken. The sample was centrifuged at 14,000 r.p.m. for 10 min, and aspirate the supernatant into a new tube. This purification step was repeated twice. Finally, DNA was precipitated with Dr. GenTLE^TM^ Precipitation Carrier (Cat#9082, Takara), sodium acetate (pH 5.2), and isopropanol.

#### Purification of the DLO Hi-C DNA fragments after MmeI digestion

Dissolved DNA was digested with 0.1 units/μl *MmeI* (Cat#R0637S, NEB) and 1.6 mM S-adenosyl-methionine (SAM) (Cat#B9003S, NEB) at 37 °C for 1 h. Then, a native polyacrylamide gel electrophoresis (PAGE) gel was applied to separate the digested DNA. The excised specific DLO Hi-C DNA fragments were subsequently placed into a 0.6-ml tube with a pierced bottom. The excised gel slices were shredded by centrifuging at 14,000 r.p.m. for 10 min. After adding TE buffer, the tubes were stored mixture at −80 °C for 20 min and were subsequently incubated at 37 °C for 2 h with simultaneous rotation at 15 r.p.m. All DNA was collected using a 2-ml Costar^®^ Spin-X^®^ tube filter (Cat#8160, Corning). Lastly, all eluate was further precipitated with Dr. GenTLE^TM^ Precipitation Carrier (Cat#9082, Takara), sodium acetate (pH 5.2), and isopropanol.

#### The library preparation of the Illumina sequencing

To ligate the Illumina sequencing adapters to the DLO Hi-C DNA fragments, PE-adaptor1, and PE-adaptor2 were added to the DLO Hi-C DNA fragments using T4 DNA ligase (Cat#15224025, Thermo Fisher Scientific) reaction mixture. After incubation at 16 °C for 30 min, AMPure XP beads (NEB) were applied to purify all DNA fragments. The eluted DNA was further repaired by PreCR Repair Mix (Cat#M0309L, NEB) by incubation at 37 °C for 20 min. Lastly, 5 μl of repaired DNA was used as a template to amplify for fewer than 13 cycles.

### Data analysis

#### Hi-C Processing

We used the *Macaca mulatta* genome (Mmul_8.0.1) as a reference genome. The DLO Hi-C tool [76] was applied to process the Hi-C data. This tool pipeline begins with raw sequencing reads and completes the following four main steps: pre-processing of raw sequencing reads, sequencing reads alignment and filtering, noise reduction and paired-end reads classification, and interaction visualization.

#### Normalization

To analyze chromosomal architecture between the saline and morphine-treated cortical neurons, a comprehensive normalization method of ICE was applied to remove systematic biases [77], including the distance between restriction sites, the GC content of trimmed ligation junctions and sequence uniqueness, and mappability at a megabase resolution. Normalized contact matrices are produced at all resolutions using the ICE approach.

#### A/B compartment identification

A/B compartments were identified as described previously [78]. To determine compartment type (active A compartment or inactive B compartment) and compartmentalization strength, a distance-dependent Hi-C contact matrix (expected data matrix) was generated, followed by computing the observed/expected (O/E) matrix, across the entire genome. Then, Pearson correlation matrices were computed using the Pearsons tool by calculating Pearson correlation values between all pairs of rows and columns in the O/E matrix. The Pearson correlation matrix was subsequently subjected to principal component analysis (PCA). According to the recommended approach [79], the eigenvector of the top three principal components (PC1, PC2, and PC3) was checked. The eigenvector showing the most aligned with the largest absolute value, GC content, or gene density was used to define the A/B compartment type. Positive eigenvector value enriches with active A compartment (gene–rich regions) and negative eigenvalue enriches with inactive B compartment (gene–poor regions). The eigenvector decomposition of the 1 Mb interaction matrices was calculated through the Hiclib package.

#### Compartment strength

To determine the effect of morphine on compartment segregation, we quantified the strength of compartmentalization according to the previously reported methods [42,80]. In brief, we re-ordered each column and row of the O/E matrices according to the value of the eigenvector, which was aligned in ascending order from left and top to right and bottom, respectively. Bins representing active A compartments and inactive B compartments were thus moved to the lower– right and upper–left corners, respectively. The saddle plots were obtained by aggregating bins across the entire genome into 50 sections. The compartment strength of each chromosome was determined as the following formula: compartment strength = (median (20% strongest AA) + median (20% strongest BB))/ (median (20% strongest AB) + median (20% strongest BA)). The value in the middle of the saddle plots was the mean compartment strength of all chromosomes.

#### TAD boundary calling

The whole genome was split into 40 kb windows and the interaction frequencies within 2 Mb upstream and downstream of each window were then compared. The directionality index was applied to determine the TAD boundary as previously reported [26]. A region was marked as a TAD border if it was between two adjacent boundaries and was shorter than 400 kb. All the intervals in the saline and morphine groups were separately determined. If a region overlapped between two groups, it was identified as an overlapping boundary.

#### Loop creation and APA

For loop analysis, the pipeline’s HiCCUPS in Juicer was applied for the discovery of locally enriched peaks [33,34]. In brief, the hic format files with the variable resolutions (2.5 Mb, 1 Mb, 500 kb, 250 kb, 100 kb, 50 kb, 25 kb, 10 kb, and 5 kb) were produced by Juicer. Then, through HiCCUPS with default parameters at resolutions of 5 kb and 10 kb, all the locally enriched peaks were identified. Furthermore, the HiCCUPSDiff in Juicer tool was applied to further analyze all differential loops determined by HiCCUPS.

We produced APA plots and linked scores to assess the quality of called loops. These analyses aggregate the local background, the signal of pixels of loops, and the pixels surrounding loops. For each loop in a given set of loops, normalized contact frequencies were calculated for the loop representation pixel and for pixels within 10 bins in both the x and y directions. To normalize for loops at various distances, each pixel was divided by the expected normalized interaction frequency at that distance to provide an observed over expected ratio. At each position in the matrix, the median observed over expected ratios was calculated and further plotted as a heatmap. The median value of the nine pixels in the lower right section of the APA plot was divided by the value of the center pixel to calculate APA scores.

#### Alignment rate calculation

The locations of restriction enzyme sites were determined using HiCNorm [81] scripts, and BEDtools [82] was utilized to produce upstream and downstream reads with predefined lengths. Then, all reads were aligned to the *Macaca mulatta* genome (Mmul_8.0.1) with the Burrows-Wheeler-Alignment (BWA) package [83]. The ratio of unique mapping reads was calculated using those reads with MAPQ quality scores greater than 20.

### ChIP-seq peak enrichment analysis

ChIP-seq signals and peaks of CTCF, DNase, and H3K27ac of rhesus macaque (GSE163177 and GSE67978) were obtained from open access database (https://www.ncbi.nlm.nih.gov/geo/query/acc.cgi?acc=GSE163177 and https://www.ncbi.nlm.nih.gov/geo/query/acc.cgi?acc=gse67978) [28,29]. TAD boundaries and chromatin loop anchors are defined with 10k width. We performed a hypergeometric test to estimate the significance probability for enrichment analysis. The odds ratio and 95% confidence interval were calculated with R package named fmsb.

### RNA-seq experimental procedures

Total RNA was extracted from sorted cortical neurons using an AxyPrepTM Multisource Total RNA Miniprep kit (Cat#AP-MN-MS-RNA-50, Axygen) according to the manufacturer’s instructions. Three replicates were used for both groups. Poly-adenylated transcripts were isolated using the NEBNext^®^ Polya mRNA magnetic isolation module in accordance with the instructions. A VAHTS stranded mRNA-seq library prep kit for Illumina^®^ (Cat#NR602, Vazyme) was then used to generate the cDNA libraries. The constructed cDNA libraries were sequenced as 150 bp paired-end reads with an Illumina HiSeq X Ten sequencer instrument. The sequenced raw data in FASTQ format were filtered out adapter, reads containing ploy-N and low-quality reads from raw data. The acquired clean reads with high quality were used for further downstream analyses.

All clean reads were aligned to a *Macaca mulatta* genome (Mmul_8.0.1). The mapped reads of each sample were subsequently assembled using StringTie with default settings. Then, all transcriptomes from both groups were merged to re-construct a comprehensive transcriptome using Perl scripts. The expression level of all transcripts and the differentially expressed transcripts in the final generated transcriptome were calculated by StringTie and DESeq2 1.18.1, respectively. The expression of each differentially expressed gene in both groups was normalized with Z-score from their FPKM values according to the formula (x – m)/s. x: the FPKM value of a given DEG in saline or morphine treatment; m: the mean of FPKM values of the corresponding DEG in both groups; s: the standard deviation.

GO analysis was performed with DAVID v6.8 Functional Annotation Tool [84]. Protein-protein interaction networks are constructed from all DEGs conjugated with differential loops using Cytoscape with 3.2.1ClueGO/CluePedia plugin [85].

### GSEA

GSEA (https://www.gsea-msigdb.org/gsea/index.jsp) [86] was applied to identify altered gene sets between the saline and morphine. Before running GSEA, the normalization of the raw read counts of protein-coding genes in the switching compartments was performed by “DESeq2”. Then, the GSEA desktop application (version 4.2.3) was conducted to enrich the altered gene sets in response to morphine administration. As gene sets, we used the reactome subset of canonical pathways (CP) (c2.cp.reactome.v7.5.1.symbols.gmt) from the molecular signatures database (MSigDB) as the reference gene sets. The number of permutations was set at 1000. The nominal *P* < 0.01 and FDR < 0.25 were considered statistically significant. A positive NES indicates enrichment in the morphine, while a negative NES indicates enrichment in the saline. The enrichment score of a single gene set is estimated by nominal *P*.

### RNA isolation for RT-qPCR

Total mRNA was directly extracted from the frozen cerebral cortex of the rhesus monkey using the AxyPrepTM Multisource RNA Miniprep kit (Cat#AP-MN-MS-RNA-50, Axygen) according to the manufacturer’s instructions. Single-stranded cDNA was reverse transcribed from extracted mRNA with a PrimeScriptTM RT reagent kit with gDNA Eraser (Cat#RR047A, Takara). Quantitative PCR reactions were performed with PowerUp^TM^ SYBR^TM^ Green Master Mix (Cat#A25742, Thermo Fisher Scientific) in QuantStudio 1 Real-Time PCR System (QS-1) (Thermo Fisher Scientific, Waltham, MA). *Macaca mulatta GAPDH* was used as a reference control. Changes in expression levels were calculated with the 2^−ΔΔCt^ method. All data were presented as mean ± SD. Differences between the saline and morphine were measured by GraphPad Prism 9 software using Student’s t-test. The *P* < 0.05 was considered statistically significant (\**P* < 0.05, ** *P* < 0.01, *** *P* < 0.001). Primers used in this study were listed in Table S14.

## Ethical statement

All monkey experiments were carried out in accordance with the guidance for the Care and Use of Laboratory Animals and with approval from the Institutional Animal Care and Use Committee (IACUC, Approval No. 2019305A) of West China Hospital, Sichuan University. All efforts were made to minimize the suffering of the animals.

## Data availability

The raw sequence data reported in this paper have been deposited in the Genome Sequence Archive (GSA) in the National Genomics Data Center, Beijing Institute of Genomics, Chinese Academy of Sciences / China National Center for Bioinformation (GSA: PRJCA012908), which is publicly accessible for reviewers at https://ngdc.cncb.ac.cn/gsa/s/7opAGr51.

Additionally, the raw sequence data generated and/or analyzed during the current study are also available in the [GEO: under accession number GSE196212] repository, https://www.ncbi.nlm.nih.gov/geo/query/acc.cgi?acc=GSE196212.

## CRediT author statement

**Liang Wang**: Methodology, Software, Validation, Formal analysis, Writing-original draft, Writing - review & editing. **Xiaojie Wang**: Methodology, Software. **Chunqi Liu**: Investigation, Formal analysis. **Wei Xu**: Investigation. **Weihong Kuang**: Resources. **Qian Bu**: Software, Investigation. **Hongchun Li**: Methodology. **Ying Zhao**: Resources. **Linhong Jiang**: Methodology. **Yaxing Chen**: Software, Resources. **Feng Qin**: Resources. **Shu Li**: Methodology. **Qinfan Wei**: Data curation. **Xiaocong Liu**: Formal analysis. **Bin Liu**: Project administration. **Yuanyuan Chen**: Methodology, Investigation. **Yanping Dai**: Project administration. **Hongbo Wang**: Conceptualization, Resources. **Jingwei Tian**: Conceptualization, Resources. **Gang Cao**: Software, Validation, and Formal analysis. **Yinglan Zhao**: Supervision, Writing - review & editing. **Xiaobo Cen**: Conceptualization, Investigation, Supervision, Review & editing, Funding acquisition. All authors have read and approved the final manuscript.

## Acknowledgments

We appreciate a lot the staff at Wuhan GeneCreate Biological Engineering Co., Ltd. (Wuhan, China) for technical assistance and advice on experiments. Our sincere thanks give to Dr. Tianming Wu from Professor Stephen Daltond’s lab for his critical suggestion on experimental design and professional advice on the analysis of Hi-C data. Lastly, we deeply acknowledge Dr. Xiaonan Liu from the University of Helsinki for his nice help on the analysis of RNA-seq data. Thanks a lot for Dr. Hongxia Zhao from the University of Helsinki for her nice help in grammar revision.

We appreciate a lot for financial support from the National Natural Science Foundation of China (Grant Nos. 82071494, 81871043, 32000719, 81272459), 1·3·5 Project for Disciplines of Excellence, West China Hospital, Sichuan University (Grant No. ZYGD18024), the China Postdoctoral Science Foundation (Grant No. 2021M702362), the Post-Doctor Research Project, West China Hospital, Sichuan University (Grant No. 2020HXBH010), Sichuan Science and Technology Program (Grant No. 23NSFSC2884), Science, Technology and Innovation Commission of Shenzhen Municipality (Grant No. ZDSYS20190902093601675).

## Supplementary material

**Figure S1.**
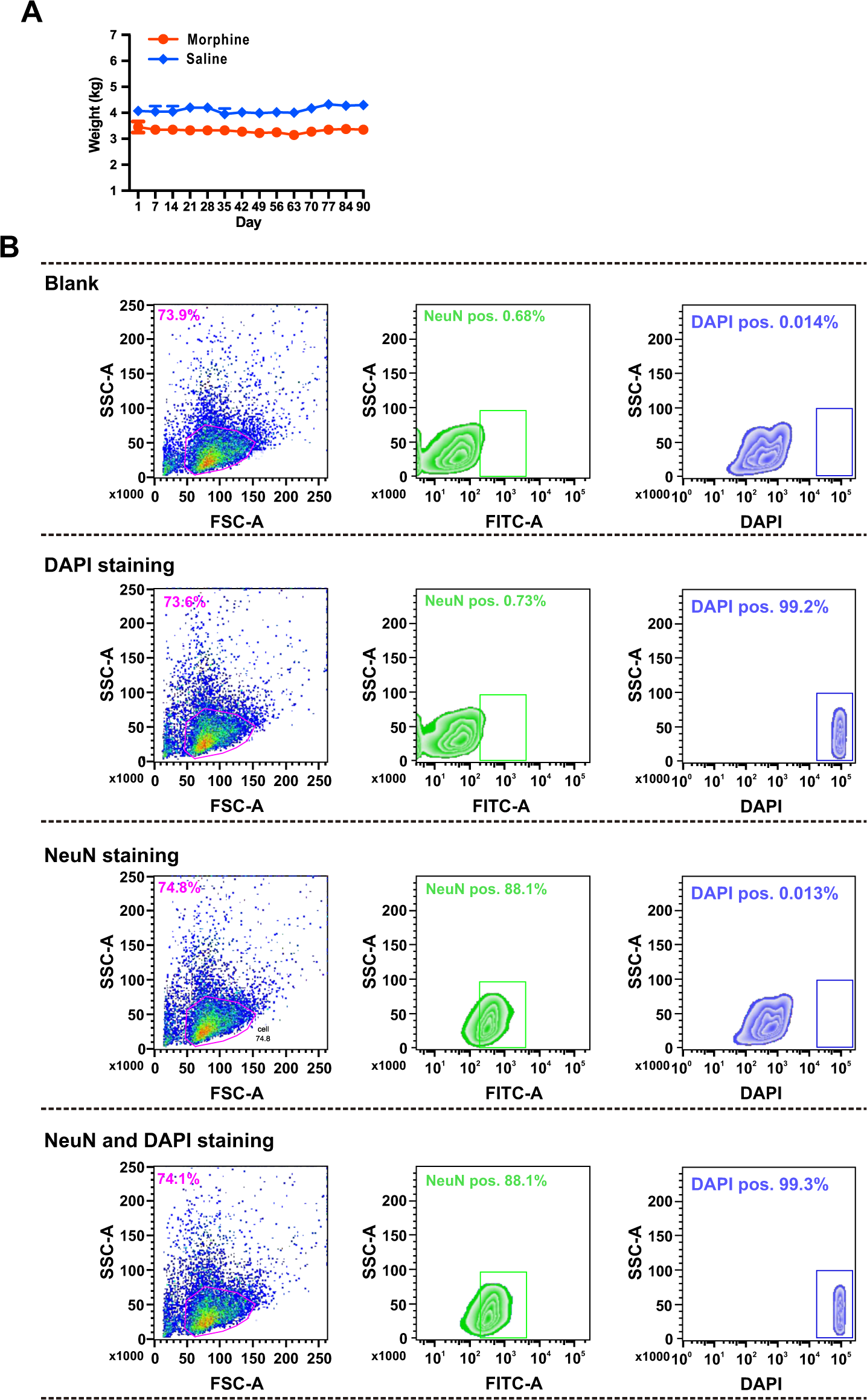
The bodyweight of the monkey and gating strategy for mature neuron cell subsets. **A.** The line plot displays the bodyweight of the rhesus monkey during 90 days of morphine (red) or saline (blue) treatment. **B.** Example flow cytometric data of unstained cells, DAPI-positive, NeuN-positive, NeuN- and DAPI-positive cells from non-human primate cortical neurons. FACS scatter plotting cell granularity based on the FSC-A on the x-axis versus SSC-A on the x-axis. FSC-A, forward scatter A; SSC-A, side scatter A.

**Figure S2.**
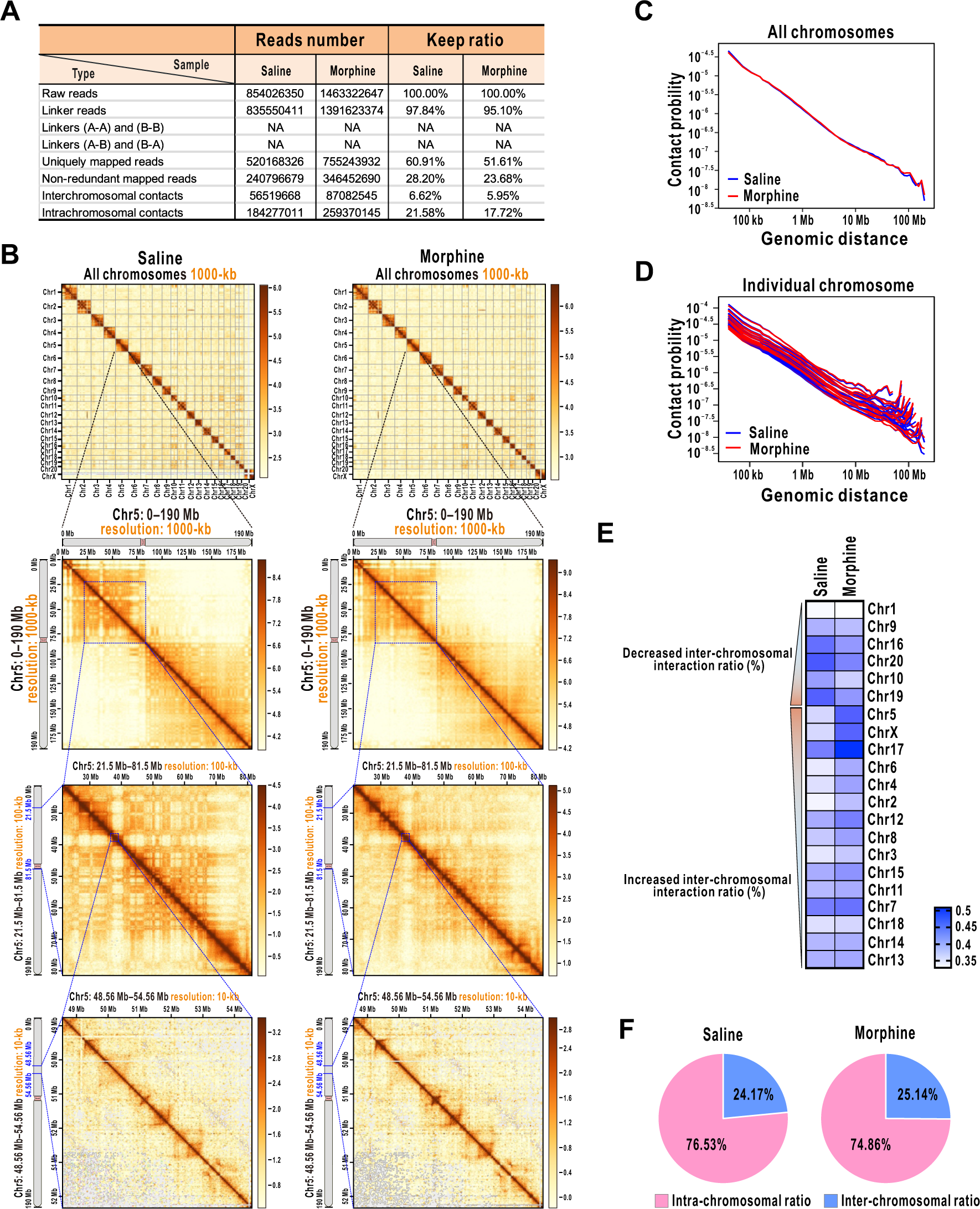
Long-term morphine administration alters chromatin spatial organization of cortical neurons. **A.** Parameters used for Hi-C analysis. **B.** Heatmaps show normalized DLO Hi-C interaction frequencies at different resolutions in the saline and morphine groups. 1000 kb for the entire genome and Chr5; 100 kb for the selected area of Chr5 (Chr5: 21.5 Mb–81.5 Mb), and 10 kb for a magnified view between 21.5 Mb–81.5 Mb of Chr5 (Chr5: 48.56 Mb–54.56 Mb). **C.** Genome– wide interaction frequency as a function of average genomic distance for saline (blue) and morphine (red). **D.** The interaction frequency of each chromosome as a function of average genomic distance for saline (blue) and morphine (red). **E.** Heatmap of trans-interaction ratio in each chromosome. Decreased and increased trans-interaction ratios are separately displayed in the up and down parts. **F.** The pie charts show the ratio of intra- and inter-chromosomal interaction.

**Figure S3.**
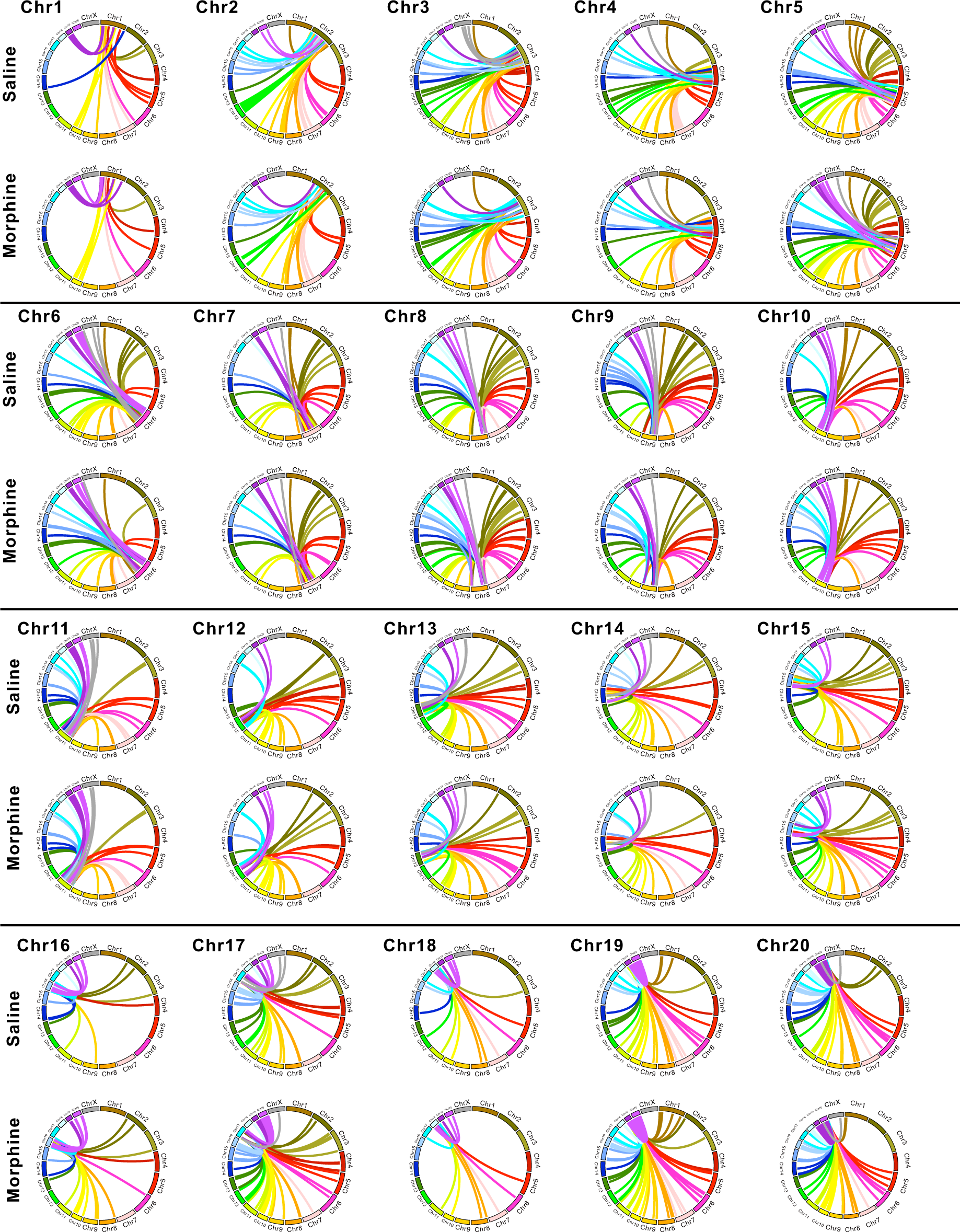
Circos plots of inter-chromosomal interactions of a single chromosome with the other chromosomes. Through investigating the first 1000 inter-chromosomal interactions, the interactions of the chromosome with the rest of the chromosomes are shown with different colors.

**Figure S4.**
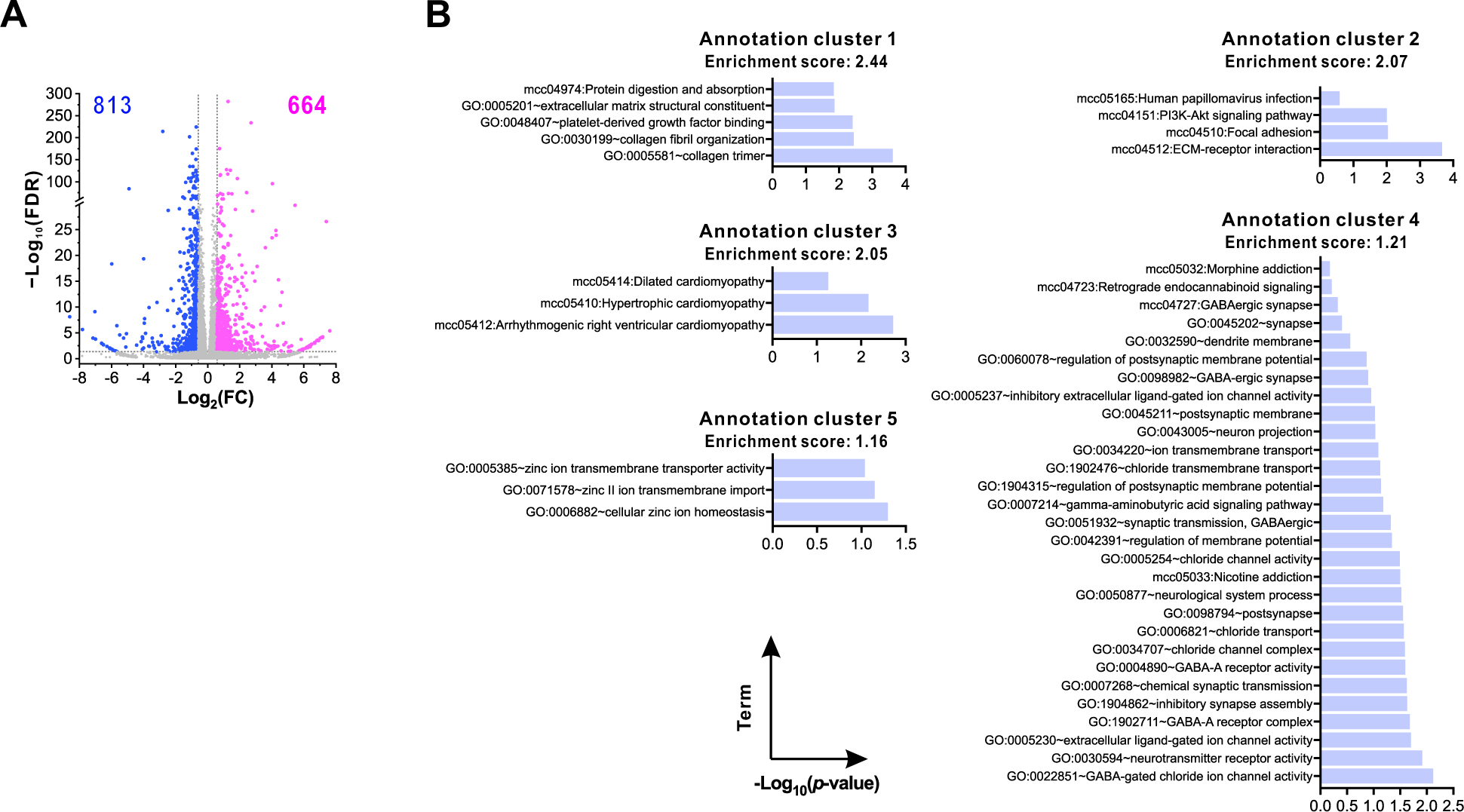
Functional analysis of all significantly changed genes in RNA-seq following morphine treatment. **A.** Volcano plot shows all significant up- and down-regulated genes (FC > 1.5 and FDR < 0.05) based on RNA-seq. **B.** The bar chart shows the top 5 enriched functional clusters of all significantly changed genes using the DAVID functional annotation tool.

**Figure S5.**
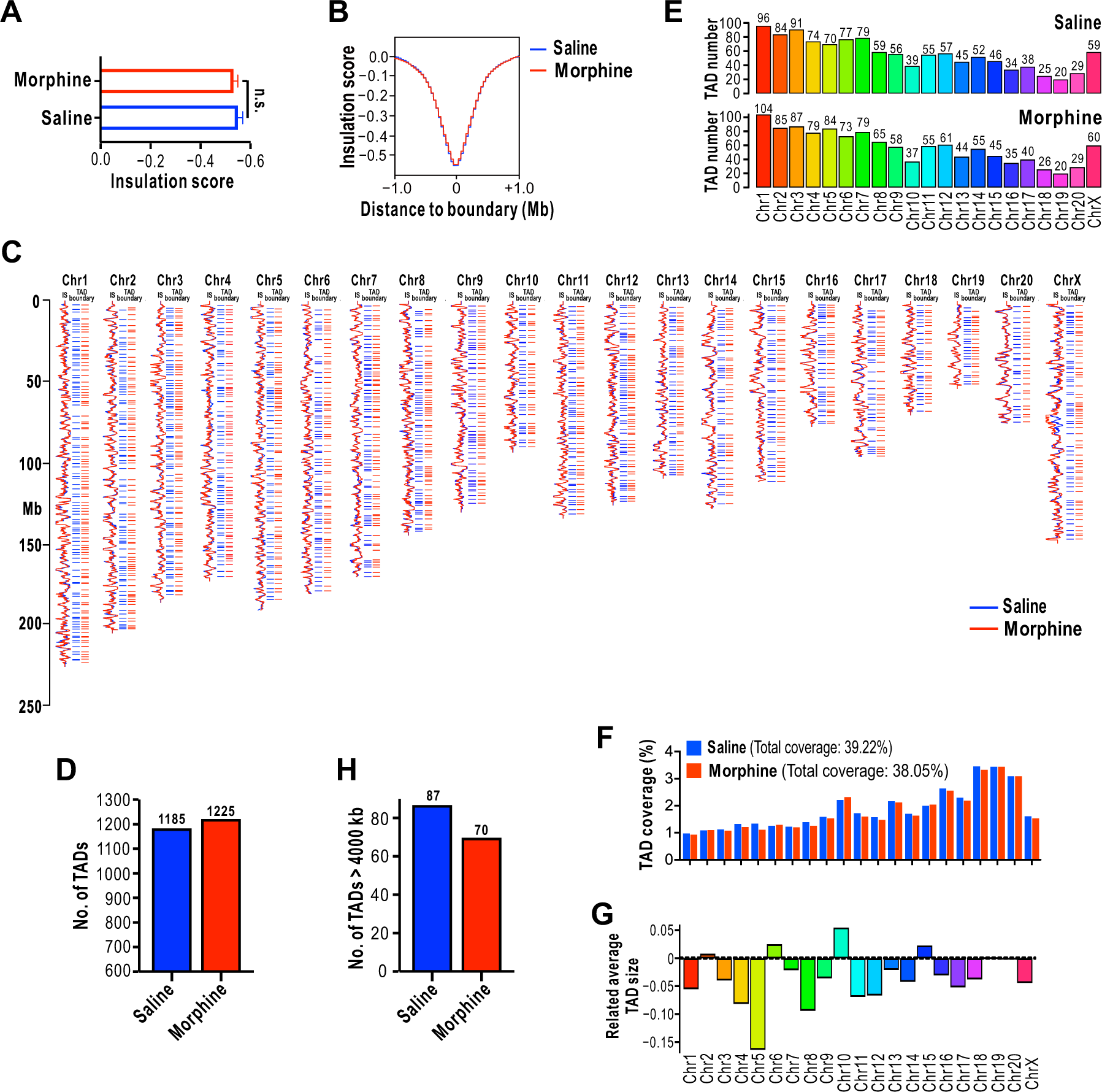
Morphine alters the TAD and TAD boundary. **A.** Bar graph shows the average insulation score of all identified TAD boundaries. **B.** Meta-region plots of insulation scores centering around ±1 Mb TAD boundaries in the saline (blue) and morphine groups (red). **C.** Snapshot of insulation score curves and distribution of identified TAD boundaries for all chromosomes in the saline (blue curve for insulation score and blue line for TAD boundaries) and morphine groups (red curve for insulation score and red line for TAD boundaries). **D.** Bar graph displays the total number of identified TADs across the entire genome. **E.** Bar graph displays the total number of identified TADs in each chromosome. **F.** Bar graph presents the percentage of TAD coverage in each chromosome. **G.** Bar graph shows the related change in the average TAD size of the morphine group in comparison to the saline group. **H.** Bar graph shows the total number of TADs with a size above 4000 kb in the saline and morphine groups.

**Figure S6.**
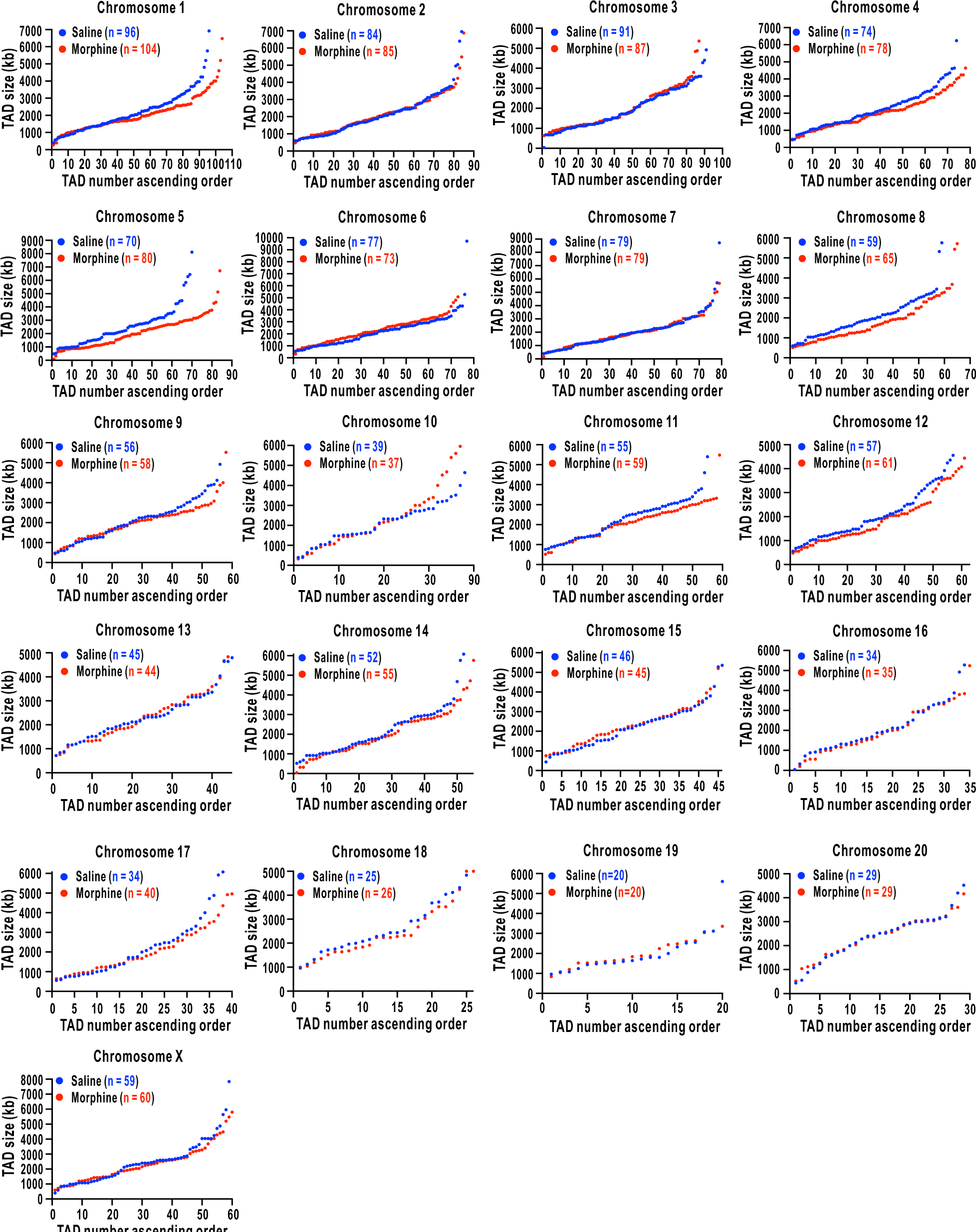
Scatter plots display the TAD size of individual chromosomes. All TADs identified in the saline (blue dot) and morphine group (red dot) were arrayed based on the size of the TADs. The total number of TADs is shown in the upper left corner of each scatter plot.

**Figure S7.**
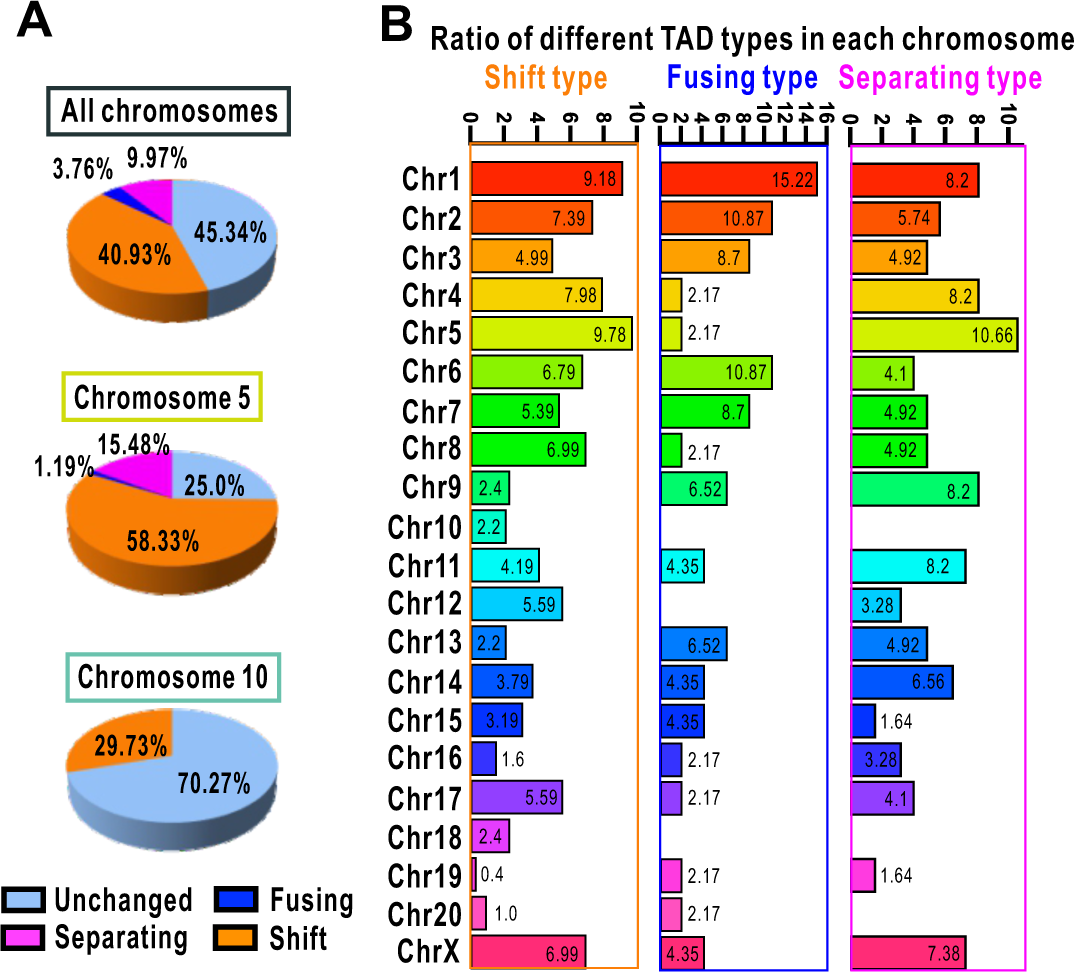
Morphine causes three types of alteration in chromatin topological structure. **A.** Pie chart shows the percentage of unchanged TADs and changed TADs across the entire genome and two additional chromosomes (Chr5 and Chr10) caused by morphine. The unchanged TAD is presented as light blue color; the changed TADs with shift, fusing, and separating types are presented as orange, dark blue, and magenta color, respectively. **B.** Bar graph displays the percentage of each TAD type in each chromosome. The different colors represent individual chromosomes, and the value at the top of the bar represents the ratio of changed TAD numbers in a single chromosome among altered TADs in all chromosomes.

**Figure S8.**
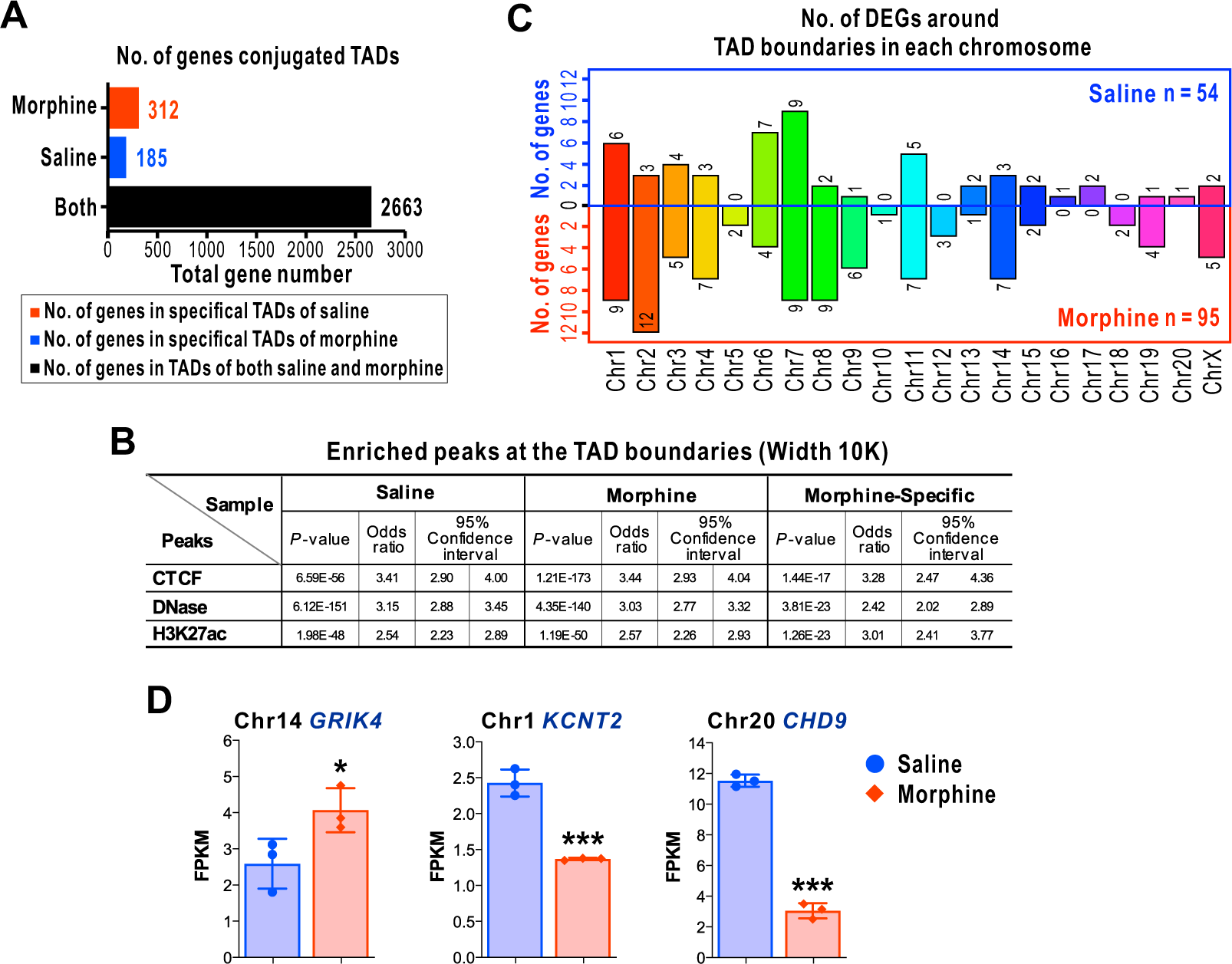
The number of changed genes correlating with TADs. **A.** Bar graph shows the number of genes around TAD boundaries identified in both groups (black). The specific TAD boundaries in the saline and morphine group are displayed in blue and red colors, respectively. **B.** The enrichment results of TAD boundaries in CTCF, DNase, and H3K27ac ChIP- seq peaks. **C.** Bar graph displays the number of significantly changed genes around specific TAD boundaries in each chromosome. **D.** Bar graph shows the FPKM value of a given DEG. Data are presented as mean ± SD and analyzed using unpaired *t* test, n = 3; \**P* < 0.05,\*\*\**P* < 0.001. Compared to the saline.

**Figure S9.**
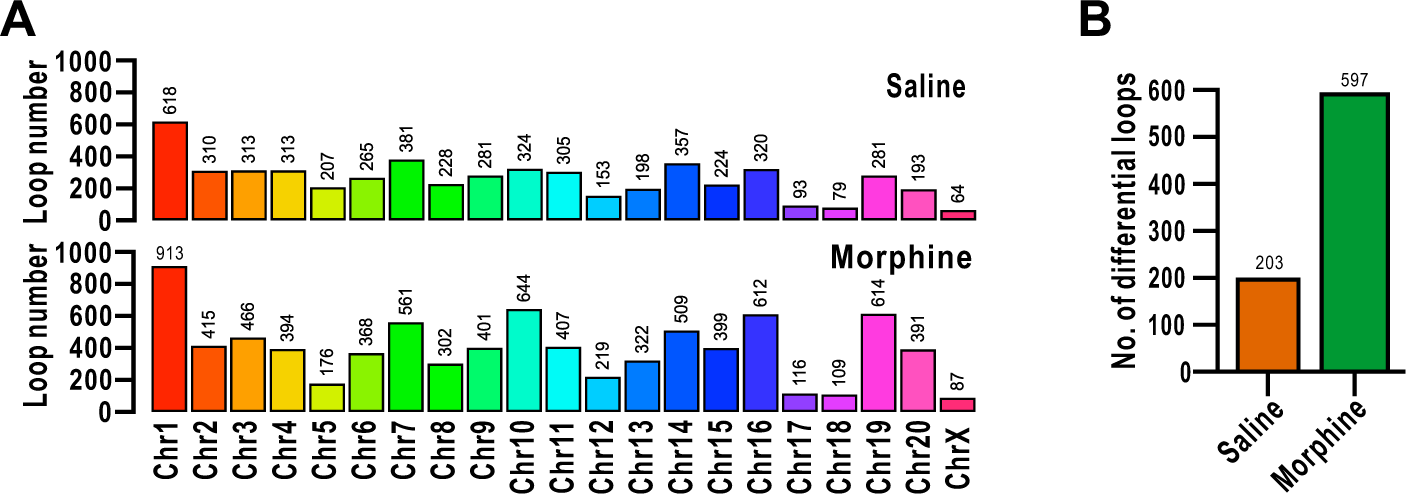
Morphine alters chromatin looping architecture. **A.** Bar graph displays the total number of loops in each chromosome. **B.** Bar graph shows the number of differential loops in the saline and morphine groups.

**Figure S10.**
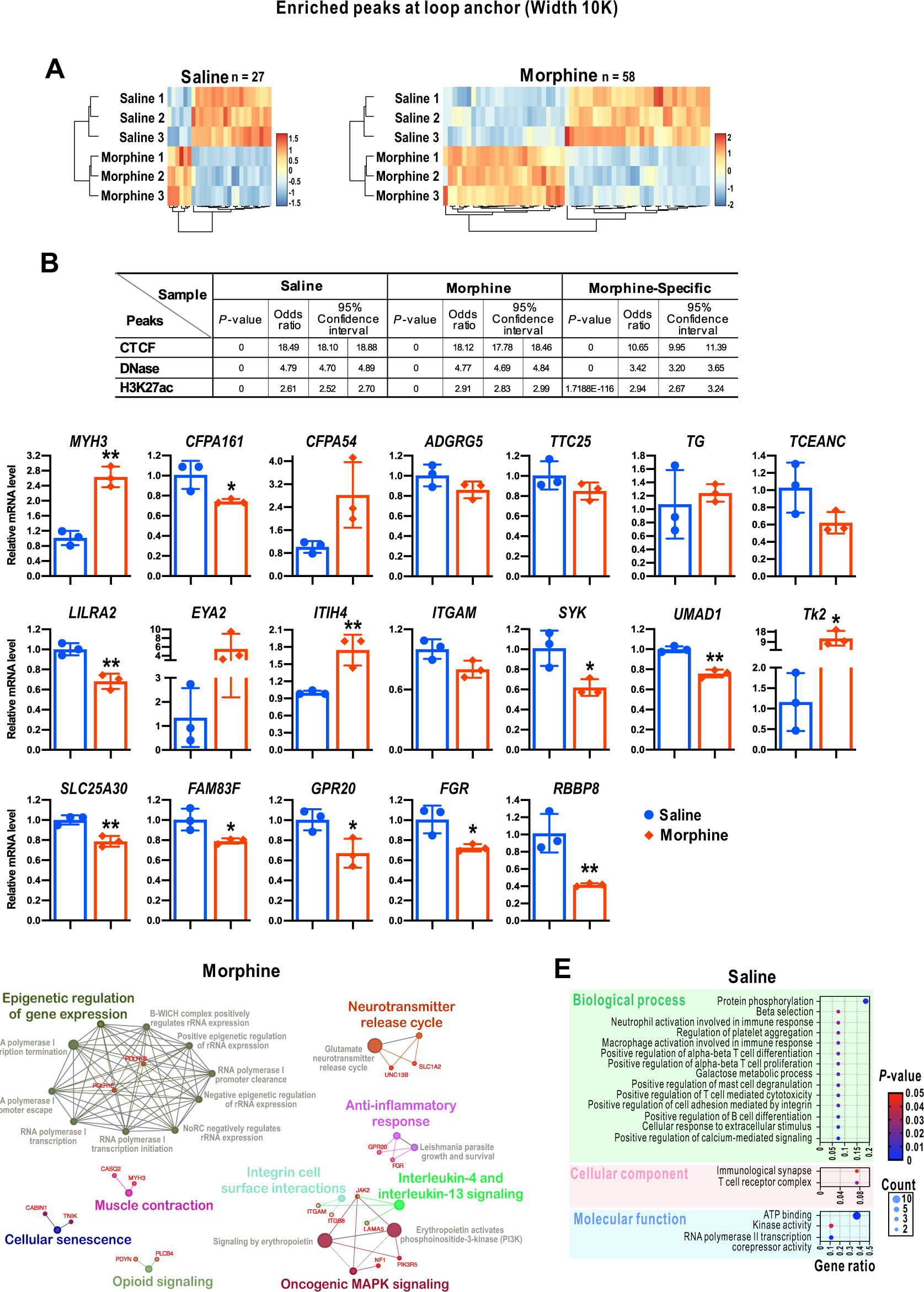
Morphine modifies cell signaling pathways through altered looping architecture. **A.** Heatmap of hierarchical clustering of DEGs linking to significantly altered looping. Each row represents an experimental treatment (Saline 1, Saline 2, Saline 3; Morphine 1, Morphine 2, Morphine 3) and each column represents a screened DEG. Orange means up-regulation and light blue means down-regulation. **B.** The enrichment results of loop anchors in CTCF, DNase, and H3K27ac ChIP-seq peaks. **C.** Bar graphs display the transcriptional levels of genes acquired from differential loops. Data from three replicates (n = 3) for each group were used for statistical analysis. All data are presented as mean ± SD and analyzed using unpaired *t* test, n = 3; \**P* < 0.05,\*\**P* < 0.01. Compared to the saline. **D.** Network enrichment analysis of DEGs in the morphine group. GO gene enrichment profiling in morphine treatment is visualized. The color code of nodes corresponds to the functional group to which they belong. Bold–colored characters signify the most essential functional terms which define the pathways within each class. Red- colored characters represent the relevant genes in each functional term. Each node constitutes a precise term. **E.** Bubble chart shows the significantly changed terms in the saline group based on GO enrichment analysis. Three aspects of GO enrichment analysis are presented. The x-axis represents the percentage of the DEGs in one aspect, and the y-axis represents the name of each cellular process. The *P* is shown in different colors. blue: low *P* value; red: high *P* value. Compared to the saline. The size of the bubble area shows the genes of DEGs that belong to one cellular process.

**Figure S11.**
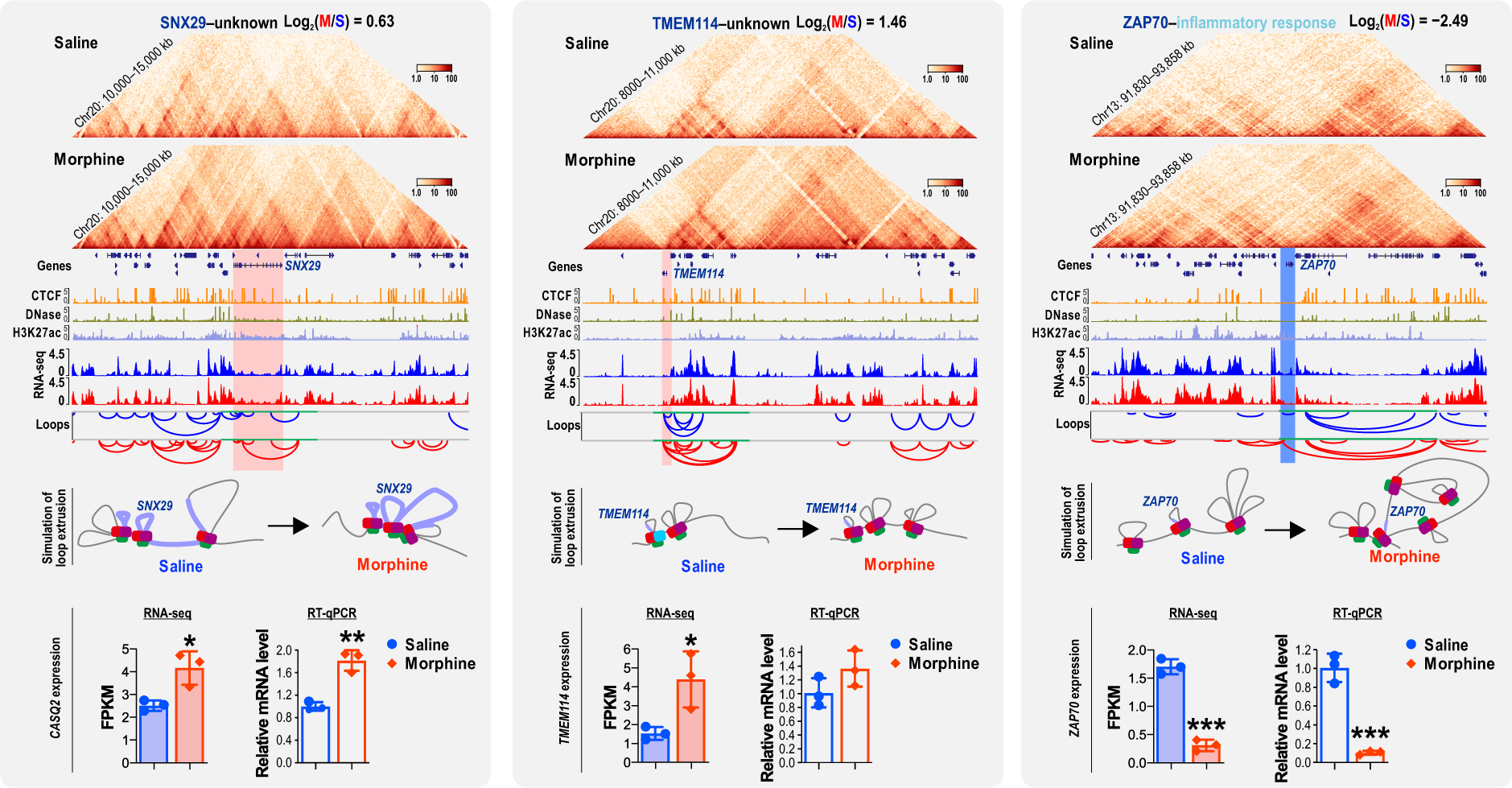
Representative genes modified by the change of chromatin loop architecture induced by morphine. The up-regulated *SNX29* and *TMEM114* as well as down-regulated *ZAP70* separately as exemplified to demonstrate that morphine evidently modulates the loop architecture, which links to the changed transcriptional activities of genes. Top two rows: Hi-C contact maps were rotated 45^°^ so that the main diagonal is horizontal. IGV screenshots of CTCF, DNase, and H3K27ac ChIP- seq peaks are presented with different colors. The location of DEGs linking to the differential loop at linear genome is marked with light orange color and light purple in simulated loop extrusion. The region for simulation of loop extrusion is covered with the green line at x-axis of loop calls. For the bar graphs of RNA-seq and RT-qPCR in the last row, data from three replicates (n = 3) for each group were used for statistical analysis. All data are presented as mean ± SD and analyzed using unpaired *t* test, n = 3, \**P* < 0.05,\*\**P* < 0.01, \*\*\**P* < 0.001.

**Table S1 Cumulative number of spontaneous withdrawal signs observed in rhesus monkeys at 30, 60, 90, 120, and 150 min, respectively, after morphine deprivation for 14 h**

**Table S2 Percentage of intra-chromosomal interaction in the saline and morphine groups**

**Table S3 Percentage of inter-chromosomal interaction in the saline and morphine groups**

**Table S4 Compartment analysis in the saline group**

**Table S5 Compartment analysis in the morphine group**

**Table S6 Change of genes involved in the A to B compartment switching after morphine administration**

**Table S7 Change of genes involved in the B to A compartment switching after morphine administration**

**Table S8 The size of all identified TADs (kb) in each chromosome**

**Table S9 Change of genes in conjugated TAD analysis**

**Table S10 Differential loops specifically identified in the saline group**

**Table S11 Differential loops specifically identified in the morphine group**

**Table S12 Differential loops and differential genes specifically identified in the saline group**

**Table S13 Differential loops and differential genes specifically identified in the morphine group**

**Table S14 Primers used for RT-qPCR assay**

## References

[1] Ammon-Treiber S, Höllt V. Morphine-induced changes of gene expression in the brain. Addict Biol 2005;10:81–9.

[2] Bu Q, Yang Y, Yan G, et al. Proteomic analysis of the nucleus accumbens in rhesus monkeys of morphine dependence and withdrawal intervention. J Proteomics 2012;75:1330–42.

[3] Ziółkowska B, Korostyński M, Piechota M, Kubik J, Przewołcki R. Effects of morphine on immediate-early gene expression in the striatum of C57BL/6J and DBA/2J mice. Pharmacol Reports 2012;64:1091–104.

[4] Baidoo N, Wolter M, Holahan MR, Teale T, Winters B, Leri F. The effects of morphine withdrawal and conditioned withdrawal on memory consolidation and c-Fos expression in the central amygdala. Addict Biol 2021;26:e12909.

[5] Sivalingam M, Ogawa S, Parhar IS. Mapping of morphine-induced OPRM1 gene expression pattern in the adult zebrafish brain. Front Neuroanat 2020;14:5.

[6] Rouhani F, Khodarahmi P, Naseh V. NGF, BDNF and Arc mRNA expression in the hippocampus of rats after administration of morphine. Neurochem Res 2019;44:2139–46.

[7] Bonev B, Cavalli G. Organization and function of the 3D genome. Nat Rev Genet 2016;17:661–78.

[8] Anania C, Lupiáñez DG. Order and disorder: abnormal 3D chromatin organization in human disease. Brief Funct Genomics 2020;19:128–38.

[9] Chakraborty A, Ay F. The role of 3D genome organization in disease: From compartments to single nucleotides. Semin Cell Dev Biol 2019;90:104–13.

[10] Spielmann M, Lupiáñez DG, Mundlos S. Structural variation in the 3D genome. Nat Rev Genet 2018;19:453–67.

[11] Franke M, Ibrahim DM, Andrey G, et al. Formation of new chromatin domains determines pathogenicity of genomic duplications. Nature 2016;538:265–9.

[12] Szabo Q, Bantignies F, Cavalli G. Principles of genome folding into topologically associating domains. Sci Adv 2019;5:eaaw1668.

[13] Saberian H, Asgari Taei A, Torkaman-Boutorabi A, et al. Effect of histone acetylation on maintenance and reinstatement of morphine-induced conditioned place preference and ΔFosB expression in the nucleus accumbens and prefrontal cortex of male rats. Behav Brain Res 2021;414:113477.

[14] Barrow TM, Byun H-M, Li X, et al. The effect of morphine upon DNA methylation in ten regions of the rat brain. Epigenetics 2017;12:1038–47.

[15] Chen M, Zhu Q, Li C, et al. Chromatin architecture reorganization in murine somatic cell nuclear transfer embryos. Nat Commun 2020;11:1–14.

[16] Wise LE, Premaratne ID, Gamage TF, et al. l-theanine attenuates abstinence signs in morphine-dependent rhesus monkeys and elicits anxiolytic-like activity in mice. Pharmacol Biochem Behav 2012;103:245–52.

[17] Denker A, de Laat W. The second decade of 3C technologies: detailed insights into nuclear organization. Genes Dev 2016;30:1357–82.

[18] Eres IE, Luo K, Hsiao CJ, Blake LE, Gilad Y. Reorganization of 3D genome structure may contribute to gene regulatory evolution in primates. PLoS Genet 2019;15:e1008278.

[19] Lin D, Hong P, Zhang S, et al. Digestion-ligation-only Hi-C is an efficient and cost-effective method for chromosome conformation capture. Nat Genet 2018;50:754–63.

[20] Dunn KE, Huhn AS, Bergeria CL, Gipson CD, Weerts EM. Non-opioid neurotransmitter systems that contribute to the opioid withdrawal syndrome: a review of preclinical and human evidence. J Pharmacol Exp Ther 2019;371:422–52.

[21] Rosin LF, Crocker O, Isenhart RL, Nguyen SC, Xu Z, Joyce EF. Chromosome territory formation attenuates the translocation potential of cells. elife 2019;8:e49553.

[22] Cremer T, Cremer M. Chromosome territories. Cold Spring Harb Perspect Biol 2010;2:a003889.

[23] Rosa-Garrido M, Chapski DJ, Schmitt AD, et al. High-resolution mapping of chromatin conformation in cardiac myocytes reveals structural remodeling of the epigenome in heart failure. Circulation 2017;136:1613–25.

[24] Roy S, Wang J, Kelschenbach J, Koodie L, Martin J. Modulation of immune function by morphine: implications for susceptibility to infection. J Neuroimmune Pharmacol 2006;1:77–89.

[25] Martini L, Whistler JL. The role of mu opioid receptor desensitization and endocytosis in morphine tolerance and dependence. Curr Opin Neurobiol 2007;17:556–64.

[26] Dixon JR, Selvaraj S, Yue F, et al. Topological domains in mammalian genomes identified by analysis of chromatin interactions. Nature 2012;485:376–80.

[27] Hansen AS, Cattoglio C, Darzacq X, Tjian R. Recent evidence that TADs and chromatin loops are dynamic structures. Nucleus 2018;9:20–32.

[28] Luo X, Liu Y, Dang D, et al. 3D Genome of macaque fetal brain reveals evolutionary innovations during primate corticogenesis. Cell 2021;184:723–740.e21.

[29] Vermunt MW, Tan SC, Castelijns B, et al. Epigenomic annotation of gene regulatory alterations during evolution of the primate brain. Nat Neurosci 2016;19:494–503.

[30] Malenka RC, Bear MF. LTP and LTD. Neuron 2004;44:5–21.

[31] Sun F, Zhou K, Tian K, et al. Atrial natriuretic peptide promotes neurite outgrowth and survival of cochlear spiral ganglion neurons in vitro through NPR-A/cGMP/PKG signaling. Front Cell Dev Biol 2021;9:1407.

[32] Siahposht-Khachaki A, Ezzatpanah S, Razavi Y, Haghparast A. NMDA receptor dependent changes in c-fos and p-CREB signaling following extinction and reinstatement of morphine place preference. Neurosci Lett 2018;662:147–51.

[33] Rao SSP, Huntley MH, Durand NC, et al. A 3D map of the human genome at kilobase resolution reveals principles of chromatin looping. Cell 2014;159:1665–80.

[34] Durand NC, Shamim MS, Machol I, et al. Juicer provides a one-click system for analyzing loop-resolution Hi-C experiments. Cell Syst 2016;3:95–8.

[35] Wu G, Haw R. Functional interaction network construction and analysis for disease discovery. Methods Mol. Biol., 2017, p. 235–53.

[36] Cheng Y, Tsai R, Sung Y, et al. Melatonin regulation of transcription in the reversal of morphine tolerance: Microarray analysis of differential gene expression. Int J Mol Med 2019;43:791–806.

[37] Rosati AG, Arre AM, Platt ML, Santos LR. Rhesus monkeys show human-like changes in gaze following across the lifespan. Proc R Soc B Biol Sci 2016;283:20160376.

[38] Zheng H, Xie W. The role of 3D genome organization in development and cell differentiation. Nat Rev Mol Cell Biol 2019;20:535–50.

[39] Tang Z, Luo OJ, Li X, et al. CTCF-mediated human 3D genome architecture reveals chromatin topology for transcription. Cell 2015;163:1611–27.

[40] Dixon JR, Gorkin DU, Ren B. Chromatin domains: the unit of chromosome organization. Mol Cell 2016;62:668–80.

[41] Nuebler J, Fudenberg G, Imakaev M, Abdennur N, Mirny LA. Chromatin organization by an interplay of loop extrusion and compartmental segregation. Proc Natl Acad Sci 2018;115:E6697–706.

[42] Nora EP, Goloborodko A, Valton A-L, et al. Targeted degradation of CTCF decouples local insulation of chromosome domains from genomic compartmentalization. Cell 2017;169:930–44.

[43] Schwarzer W, Abdennur N, Goloborodko A, et al. Two independent modes of chromatin organization revealed by cohesin removal. Nature 2017;551:51–6.

[44] Quinodoz SA, Ollikainen N, Tabak B, et al. Higher-Order Inter-chromosomal Hubs Shape 3D Genome Organization in the Nucleus. Cell 2018;174:744–757.e24.

[45] Flavahan WA, Drier Y, Liau BB, et al. Insulator dysfunction and oncogene activation in IDH mutant gliomas. Nature 2016;529:110–4.

[46] Hnisz D, Weintraub AS, Day DS, et al. Activation of proto-oncogenes by disruption of chromosome neighborhoods. Science (80) 2016;351:1454–8.

[47] Delaneau O, Zazhytska M, Borel C, et al. Chromatin three-dimensional interactions mediate genetic effects on gene expression. Science (80-) 2019;364:eaat8266.

[48] Kaiser VB, Semple CA. When TADs go bad: chromatin structure and nuclear organisation in human disease. F1000Research 2017;6:314.

[49] Giorgio E, Robyr D, Spielmann M, et al. A large genomic deletion leads to enhancer adoption by the lamin B1 gene: a second path to autosomal dominant adult-onset demyelinating leukodystrophy (ADLD). Hum Mol Genet 2015;24:3143–54.

[50] McArthur E, Capra JA. Topologically associating domain boundaries that are stable across diverse cell types are evolutionarily constrained and enriched for heritability. Am J Hum Genet 2021;108:269–83.

[51] Aller MI, Pecoraro V, Paternain A V., Canals S, Lerma J. Increased Dosage of High-Affinity Kainate Receptor Gene grik4 Alters Synaptic Transmission and Reproduces Autism Spectrum Disorders Features. J Neurosci 2015;35:13619–28.

[52] Loke MF, Wei H, Yeo J, Sng B-L, Sia AT, Tan E-C. Deep sequencing analysis to identify novel and rare variants in pain-related genes in patients with acute postoperative pain and high morphine use. J Pain Res 2019;12:2755–70.

[53] Wang H, Xu J, Lazarovici P, Quirion R, Zheng W. cAMP response element-binding protein (CREB): a possible signaling molecule link in the pathophysiology of schizophrenia. Front Mol Neurosci 2018;11:255.

[54] Ou J, Zhou Y, Li C, et al. Sinomenine protects against morphine dependence through the NMDAR1/CAMKII/CREB pathway: a possible role of astrocyte-derived exosomes. Molecules 2018;23:2370.

[55] Dacher M, Nugent FS. Morphine-induced modulation of LTD at GABAergic synapses in the ventral tegmental area. Neuropharmacology 2011;61:1166–71.

[56] Nugent FS, Penick EC, Kauer JA. Opioids block long-term potentiation of inhibitory synapses. Nature 2007;446:1086–90.

[57] Madhavan A, Bonci A, Whistler JL. Opioid-induced GABA potentiation after chronic morphine attenuates the rewarding effects of opioids in the ventral tegmental area. J Neurosci 2010;30:14029–35.

[58] Hervera A, Leánez S, Pol O. The inhibition of the nitric oxide–cGMP–PKG–JNK signaling pathway avoids the development of tolerance to the local antiallodynic effects produced by morphine during neuropathic pain. Eur J Pharmacol 2012;685:42–51.

[59] Cooper TE, Chen J, Wiffen PJ, et al. Morphine for chronic neuropathic pain in adults. Cochrane Database Syst Rev 2017;2019:1.

[60] Mehrabadi S, Karimiyan SM. Morphine Tolerance Effects on Neurotransmitters and Related Receptors: Definition, Overview and Update. J Pharm Res Int 2018;23:1–11.

[61] House S. Chronic morphine potentiates the inflammatory response by disrupting interleukin-1β modulation of the hypothalamic–pituitary–adrenal axis. J Neuroimmunol 2001;118:277–85.

[62] Listos J, Łupina M, Talarek S, Mazur A, Orzelska-Górka J, Kotlińska J. The mechanisms involved in morphine addiction: an overview. Int J Mol Sci 2019;20:4302.

[63] Alexander RP, Fang G, Rozowsky J, Snyder M, Gerstein MB. Annotating non-coding regions of the genome. Nat Rev Genet 2010;11:559–71.

[64] Mirny LA, Solovei I. Keeping chromatin in the loop(s). Nat Rev Mol Cell Biol 2021;22:439–40.

[65] Kadauke S, Blobel GA. Chromatin loops in gene regulation. Biochim Biophys Acta - Gene Regul Mech 2009;1789:17–25.

[66] Nevin LM, Xiao T, Staub W, Baier H. Topoisomerase IIβ is required for lamina-specific targeting of retinal ganglion cell axons and dendrites. Development 2011;138:2457–65.

[67] Hu H, Wang H, Xiao Y, et al. Otud7b facilitates T cell activation and inflammatory responses by regulating Zap70 ubiquitination. J Exp Med 2016;213:399–414.

[68] Stern PR. Addicts Lose Plasticity. Sci Signal 2010;3:ec201–ec201.

[69] Chaves C, Gómez-Zepeda D, Auvity S, et al. Effect of subchronic intravenous morphine infusion and naloxone-precipitated morphine withdrawal on p-gp and bcrp at the rat blood– brain barrier. J Pharm Sci 2016;105:350–8.

[70] Butler M, Rafi S, Hossain W, Stephan D, Manzardo A. Whole exome sequencing in females with autism implicates novel and candidate genes. Int J Mol Sci 2015;16:1312–35.

[71] Xie K, Colgan LA, Dao MT, et al. NF1 is a direct G protein effector essential for opioid signaling to Ras in the striatum. Curr Biol 2016;26:2992–3003.

[72] Koijam AS, Chakraborty B, Mukhopadhyay K, Rajamma U, Haobam R. A single nucleotide polymorphism in OPRM1 (rs483481) and risk for heroin use disorder. J Addict Dis 2020;38:214–22.

[73] Chen J-H, Zhao Y, Khan RAW, et al. SNX29, a new susceptibility gene shared with major mental disorders in Han Chinese population. World J Biol Psychiatry 2021;22:526–34.

[74] Aceto MD, Carchman RA, Harris LS, Flora RE. Caffeine elicited withdrawal signs in morphine-dependent rhesus monkeys. Eur J Pharmacol 1978;50:203–7.

[75] Preissl S, Fang R, Huang H, et al. Single-nucleus analysis of accessible chromatin in developing mouse forebrain reveals cell-type-specific transcriptional regulation. Nat Neurosci 2018;21:432–9.

[76] Hong P, Jiang H, Xu W, et al. The DLO Hi-C tool for digestion-ligation-only Hi-C chromosome conformation capture data analysis. Genes (Basel) 2020;11:289.

[77] Imakaev M, Fudenberg G, McCord RP, et al. Iterative correction of Hi-C data reveals hallmarks of chromosome organization. Nat Methods 2012;9:999–1003.

[78] Miura H, Poonperm R, Takahashi S, Hiratani I. Practical analysis of Hi-C data: generating A/B compartment profiles. Methods Mol. Biol., 2018, p. 221–45.

[79] Lieberman-Aiden E, van Berkum NL, Williams L, et al. Comprehensive Mapping of Long-Range Interactions Reveals Folding Principles of the Human Genome. Science (80-) 2009;326:289–93.

[80] Zhang H, Emerson DJ, Gilgenast TG, et al. Chromatin structure dynamics during the mitosis-to-G1 phase transition. Nature 2019;576:158–62.

[81] Hu M, Deng K, Selvaraj S, Qin Z, Ren B, Liu JS. HiCNorm: removing biases in Hi-C data via Poisson regression. Bioinformatics 2012;28:3131–3.

[82] Quinlan AR, Hall IM. BEDTools: a flexible suite of utilities for comparing genomic features. Bioinformatics 2010;26:841–2.

[83] Li H, Durbin R. Fast and accurate short read alignment with Burrows-Wheeler transform. Bioinformatics 2009;25:1754–60.

[84] Huang DW, Sherman BT, Lempicki RA. Systematic and integrative analysis of large gene lists using DAVID bioinformatics resources. Nat Protoc 2009;4:44–57.

[85] Cox J, Mann M. MaxQuant enables high peptide identification rates, individualized p.p.b.- range mass accuracies and proteome-wide protein quantification. Nat Biotechnol 2008;26:1367–72.

[86] Subramanian A, Tamayo P, Mootha VK, et al. Gene set enrichment analysis: A knowledge-based approach for interpreting genome-wide expression profiles. Proc Natl Acad Sci 2005;102:15545–50.

